# Distinct signatures of loss of consciousness during Focal Impaired Awareness versus Focal to Bilateral Tonic Clonic seizures

**DOI:** 10.1101/2021.10.01.462586

**Authors:** Elsa Juan, Urszula Górska, Csaba Kozma, Cynthia Papantonatos, Tom Bugnon, Colin Denis, Vaclav Kremen, Greg Worrell, Aaron F. Struck, Lisa M. Bateman, Edward M. Merricks, Hal Blumenfeld, Giulio Tononi, Catherine Schevon, Melanie Boly

## Abstract

Loss of consciousness (LOC) is a hallmark of many epileptic seizures and carries risks of serious injury and sudden death. While cortical sleep-like activities accompany LOC during focal impaired awareness (FIA) seizures, the mechanisms of LOC during focal to bilateral tonic-clonic (FBTC) seizures remain unclear. Quantifying differences in markers of cortical activation and ictal recruitment between FIA and FBTC seizures may also help to understand their different consequences for clinical outcomes and to optimize neuromodulation therapies.

We quantified clinical signs of LOC and intracranial EEG (iEEG) activity during 129 FIA and 50 FBTC from 41 patients. We characterized iEEG changes both in the seizure onset zone (SOZ) and in areas remote from SOZ with a total of 3386 electrodes distributed across brain areas. First, we compared the dynamics of iEEG sleep-like activities: slow-wave activity (SWA; 1-4 Hz) and beta/delta ratio (B/D; a validated marker of cortical activation) during FIA vs. FBTC. Second, we quantified differences between FBTC and FIA for a marker validated to detect ictal cross-frequency coupling: phase-locked high-gamma (PLHG; high gamma phased locked to low frequencies) and a marker of ictal recruitment: the epileptogenicity index (i.e. the number of channels crossing an energy ratio threshold for high vs. low frequency power). Third, we assessed changes in iEEG activity preceding and accompanying behavioral generalization onset and their correlation with electromyogram (EMG) channels. In addition, we analyzed human cortical multi-unit activity recorded with Utah arrays during three FBTC.

Compared to FIA, FBTC seizures were characterized by deeper LOC and by stronger increases in SWA in parieto-occipital cortex. FBTC also displayed more widespread increases in cortical activation (B/D), ictal cross-frequency coupling (PLHG) and ictal recruitment (epileptogenicity index). Even before generalization, FBTC displayed deeper LOC; this early LOC was accompanied by a paradoxical increase in B/D in fronto-parietal cortex. Behavioral generalization coincided with complete loss of responsiveness and a subsequent increase in high-gamma in the whole brain, which was especially synchronous in deep sources and could not be explained by EMG. Similarly, multi-unit activity analysis of FBTC revealed sustained increases in cortical firing rates during and after generalization onset in areas remote from the SOZ.

Unlike during FIA, LOC during FBTC is characterized by a paradoxical increase in cortical activation and neuronal firing. These findings suggest differences in the mechanisms of ictal LOC between FIA and FBTC and may account for the more negative prognostic consequences of FBTC.

## Introduction

Epilepsy is a frequent and disabling disease. In the US, 5% of the population will have at least one seizure in their lifetime, while chronic epilepsy affects approximately 3 million adults and 470,000 children.^1^ About one third of chronic epilepsies are refractory to pharmacological treatment,^2^ and surgery to achieve seizure freedom can only be performed in a minority of these cases.^3^ Seizures that are accompanied by loss of consciousness (LOC) have especially detrimental consequences on quality of life, partly through their impact on driving limitations and on social stigmatization.^4,5^ Among all seizure types, focal to bilateral tonic-clonic (FBTC; previously called ‘secondary generalized’) seizures are the most disabling, due to a complete inability to control behavior and an increased risk of sudden death.^6–8^ Although focal impaired awareness (FIA; previously called ‘complex partial ‘) seizures can also lead to serious injuries – for example if they occur while driving – they are usually characterized by a partial impairment of responsiveness.^9,10^ Unlike FBTC, a partial recall of subjective experiences is often observed after FIA.^11,12^ Importantly, frequent FBTC also predict poorer cognitive and surgical outcomes compared to frequent FIA.^13^

In recent years, direct intracranial electroencephalography (iEEG) studies in humans showed that during FIA of temporal lobe onset, sleep-like slow-wave activity (SWA; 1-4 Hz) is seen in widespread bilateral cortical networks, while ictal activity itself is restricted to a small area surrounding the seizure onset zone (SOZ).^14^ This discovery led to the development of promising neuromodulation therapies targeting arousal centers to reverse LOC during FIA.^15,16^ While it was reported that FIA and FBTC share similar electrographic ictal onset patterns,^17^ the electrophysiological correlates of behavioral generalization and LOC during FBTC have not yet been quantified. A better understanding of the cortical dynamics driving the evolution of focal seizures towards generalization could have broad implications for preventive approaches to FBTC.

Previous studies suggested that ictal activity is limited to a small cortical area during FIA.^14^ However, the occurrence of high-frequency oscillations (HFO, >80 Hz) beyond SOZ was reported during FIA in other studies.^18^ While increased HFO has also been reported during FBTC,^19^ previous studies in small samples suggested that they may not invade the whole cortex.^17,20,21^ Importantly, increased HFO do not per se signal ictal recruitment; they can increase during deep non-REM sleep^22^ and in the ictal penumbra (cortical areas that are not actively seizing).^23^ Recently, delayed-onset ictal high gamma activity (80-150 Hz) phase-locked to lower frequencies (4-30 Hz) (“phase-locked high-gamma”, PLHG)^18^ has been shown to constitute a reliable proxy for synchronized multi-unit firing bursts in the actively seizing cortex (ictal core)^24,25^. PLHG applied to clinical iEEG recordings^26^ has also been shown to predict surgical outcomes – the current gold standard to assess localization accuracy for the epileptogenic zone – more accurately than HFOs alone.^18,27^ Thus, we here sought to quantify differences in ictal cross-frequency coupling between FIA and FBTC using PLHG. As a marker of spatial ictal recruitment, we also compared the number of channels passing a threshold of energy ratio (ER) between high and low frequencies - which is at the basis of the computation of the epileptogenicity index (EI)^28^ – between FIA and FBTC. The EI detects the ordering through which each channel crosses the threshold for ictal recruitment to define SOZ; it can successfully predict surgical outcome.^29^

Here we aimed to address four main questions: (1) whether LOC during FBTC (as compared to FIA) is accompanied by sleep-like activities, (2) whether ictal recruitment is widespread during FBTC, (3) what are the iEEG signatures of behavioral generalization, and (4) how does neuronal firing change during FBTC in areas remote from the SOZ.

By comparing the temporal and spatial evolution of iEEG sleep-like activities and ictal rhythms in FBTC and FIA across 179 seizures from several academic centers, our results provide new insights about the electrocortical and neuronal firing patterns involved in secondary generalization and suggest different mechanisms for LOC during FBTC compared to FIA. The current results may have implications to understand mechanisms of LOC and differences in clinical outcomes and could have implication for preventive treatments and neuromodulation strategies.

## Material and Methods

### Datasets

Seizures were retrospectively collected from University of Wisconsin-Madison (UW-Madison) Hospital and Clinics (UWHC) medical records, from the iEEG.org online database,^30^ and from the European Epilepsy Database (EED^31^). Only seizures acquired at 400 Hz sampling rate or higher, with good quality iEEG recording, with pre- and post-electrodes implantation CT and MRI scans (or that included electrodes’ MNI coordinates) and with a reliable seizure type scoring (FBTC vs. FIA) were considered. A total of 179 seizures (50 FBTC, 129 FIA) from 41 epilepsy patients (19 female, median age 33 years old, range 14-63) implanted with intracranial electrodes were eventually included: 55 seizures from UWHC, 34 seizures from iEEG.org (mostly from the Mayo Clinic) and 90 seizures from the EED. The number of seizures per patient was lower for FBTC than for FIA (resp.1.1±1.5 and 3±3.8; *t*_(178)_=2, *p*=0.006; see Table 1). Most seizures originated from the temporal lobe (80% FBTC and 87% FIA; *p*=0.3 Fisher; see Table 1 for an exhaustive breakdown). Most seizures originated from the left hemisphere, but this was more often the case for FBTC than for FIA (57% of the FIA and 76% of the FBTC; *p*=0.02 Fisher). A comparable number of seizures arose from sleep in both seizure types (see Table 1). Considering electrographic onset patterns, most FTBC originated from rhythmic activity (48% of FBTC), while FIA often arose from low amplitude fast activity (LAFA; 40% of FIA); spiking activity was the least common onset pattern in both seizure type (20% of FBTC, 23% of FIA). No difference in electrographic onset pattern were found between FBTC and FIA (*p*=0.4, *χ*^*2*^ = 2.04). Stereo-EEG (SEEG) and subdural grids and depth electrodes (SGDE) were used to a similar extent in FIA and FBTC (see Table 1). FBTC lasted on average 121 ± 77 seconds and FIA 107 ± 69 seconds (*t*_(178)_=-1.23, *p*=0.2).

**Table 1.**
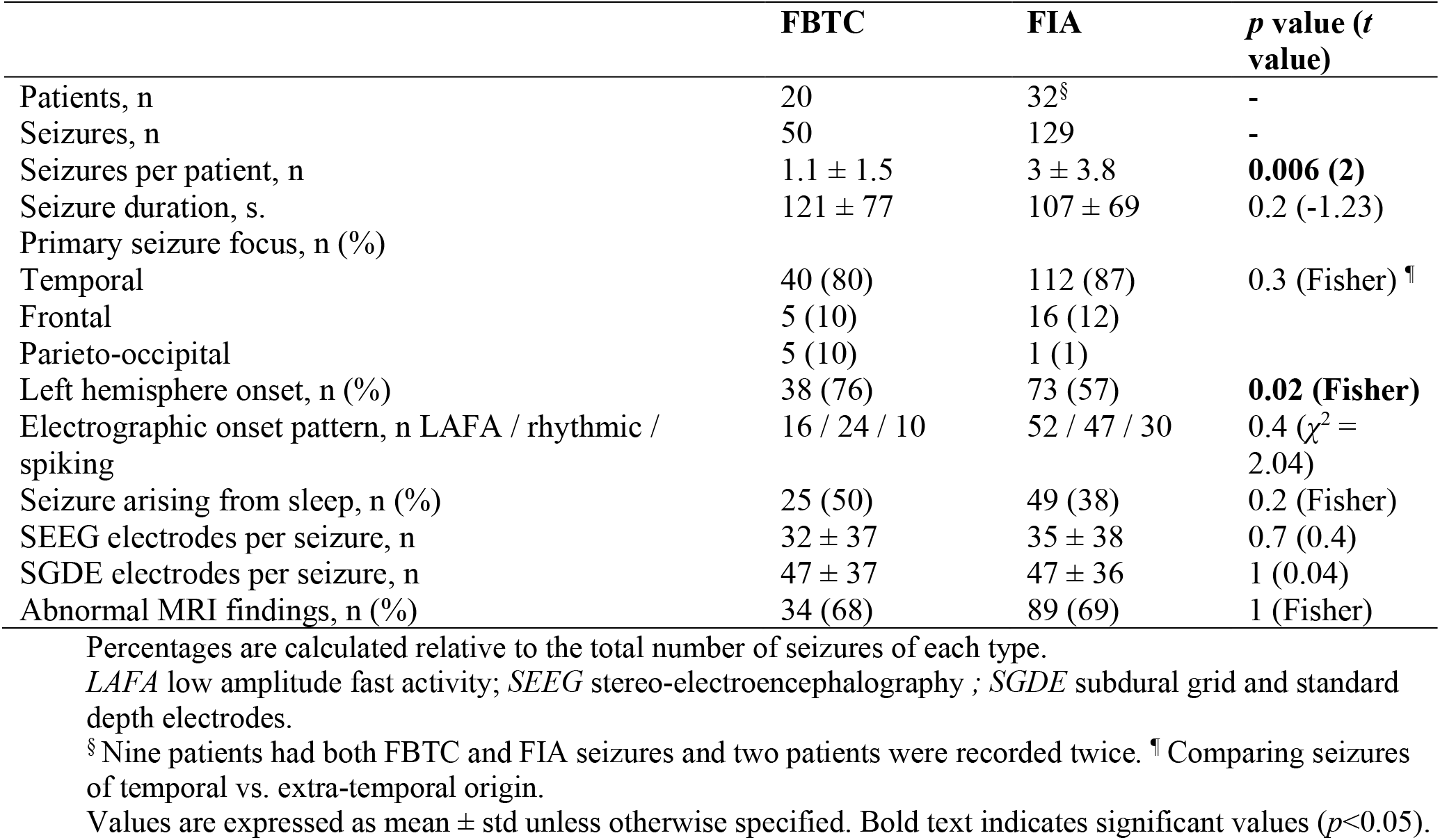
Patient’s clinical characteristics split by seizure type.

|  | FBTC | FIA | *p* value (*t* value) |
| --- | --- | --- | --- |
| Patients, n | 20 | 32<sup>§</sup> | - |
| Seizures, n | 50 | 129 | - |
| Seizures per patient, n | 1.1 ± 1.5 | 3 ± 3.8 | **0.006 (2)** |
| Seizure duration, s. | 121 ± 77 | 107 ± 69 | 0.2 (-1.23) |
| Primary seizure focus, n (%) |  |  |  |
| Temporal | 40 (80) | 112 (87) | 0.3 (Fisher) <sup>¶</sup> |
| Frontal | 5 (10) | 16 (12) |  |
| Parieto-occipital | 5 (10) | 1 (1) |  |
| Left hemisphere onset, n (%) | 38 (76) | 73 (57) | **0.02 (Fisher)** |
| Electrographic onset pattern, n LAFA / rhythmic / spiking | 16 / 24 / 10 | 52 / 47 / 30 | 0.4 ($\chi^2$ = 2.04) |
| Seizure arising from sleep, n (%) | 25 (50) | 49 (38) | 0.2 (Fisher) |
| SEEG electrodes per seizure, n | 32 ± 37 | 35 ± 38 | 0.7 (0.4) |
| SGDE electrodes per seizure, n | 47 ± 37 | 47 ± 36 | 1 (0.04) |
| Abnormal MRI findings, n (%) | 34 (68) | 89 (69) | 1 (Fisher) |

Percentages are calculated relative to the total number of seizures of each type. *LAFA* low amplitude fast activity; *SEEG* stereo-electroencephalography ; *SGDE* subdural grid and standard depth electrodes. <sup>§</sup> Nine patients had both FBTC and FIA seizures and two patients were recorded twice. <sup>¶</sup> Comparing seizures of temporal vs. extra-temporal origin. Values are expressed as mean ± std unless otherwise specified. Bold text indicates significant values (*p*<0.05).

Three additional FBTC from Columbia University from two patients with left temporal focus (two males, 28 and 29 years old) containing both iEEG and Utah microelectrodes arrays recordings were included for firing rate analysis. All procedures were approved by the institutional review board for human studies at the University of Wisconsin-Madison and at Columbia University. Informed consent was collected for the two patients prospectively enrolled at Colombia University.

### Electrographic timing, seizure categorization and behavioral scoring

The timing of seizure onset and offset was determined electrographically by a certified epileptologist (MB) for each seizure. Seizure onset time was established at the first sign of epileptic activity (e.g. heralding spike) on any channel. Seizure offset time was established when epileptic activity had ceased on all channels. The seizure onset zone was determined based on the first channels to show epileptiform activity.

Seizure classification was determined based on behavioral manifestations occurring in the ictal period. According to the 2017 ILAE guidelines,^32^ we categorized seizures as ‘focal impaired awareness’ (FIA) when response to commands or to questions was impaired or when there was amnesia at any point during the seizure (as in ^14^). ‘Focal to bilateral tonic-clonic’ seizures were recognized based on the additional occurrence of bilateral stiffening (tonic phase) followed by bilateral convulsions (clonic phase). We considered the start of the tonic phase as the onset of behavioral generalization.

When simultaneous audio and video recordings were available (e.g. in UWHC, iEEG.org and Columbia datasets), the timing of behavioral manifestations in relation to the iEEG signal was also recorded. Two main dimensions were considered: ability to follow simple commands (with scores split between verbal responsiveness and motor responsiveness), and amnesia.^14^ We also classified patient’s behavior on some of the dimensions of the Consciousness in Seizures Scale^33^ that could be consistently assessed: whether the patient was able to interact with an examiner (CSS 3), and whether the patient was aware of having a seizure while it occurred (CSS 4). Behavior was further quantified separately for the first and second half of FIA, and for the pre-generalization and the post-generalization periods of FBTC.

For seizures without video recordings (e.g. all EED seizures and some iEEG.org seizures), we relied on information provided by the database for seizures classification. These seizures were not included in analyses focusing on temporal evolution of behavior.

### Intracranial data preprocessing

Electrode localization was done semi-automatically using the iElectrodes toolbox^34^ and the FMRIB Software Library (FSL^35^). The post-implantation CT scan was aligned to the pre-implantation MRI and registered to standard MNI space. Corresponding electrodes’ voxels were then manually selected, automatically clustered and accordingly labelled (Fig.1 A1-A3). Based on the MNI coordinates identified for each electrode, probabilistic assignment to brain regions were obtained using the Talairach client.^36,37^ Contacts were then grouped in temporal, frontal and parieto-occipital areas. Following Lundstrom et al. (2019) approach,^38^ electrodes within 2.5 cm distance from the clinically identified seizure onset zone (SOZ) were further separated from other electrodes and grouped as ‘SOZ’ area. It is important to note that although most seizures originated from the temporal lobe, some seizures were of extra-temporal origin; therefore SOZ electrodes include extra-temporal locations.

**Fig. 1.**
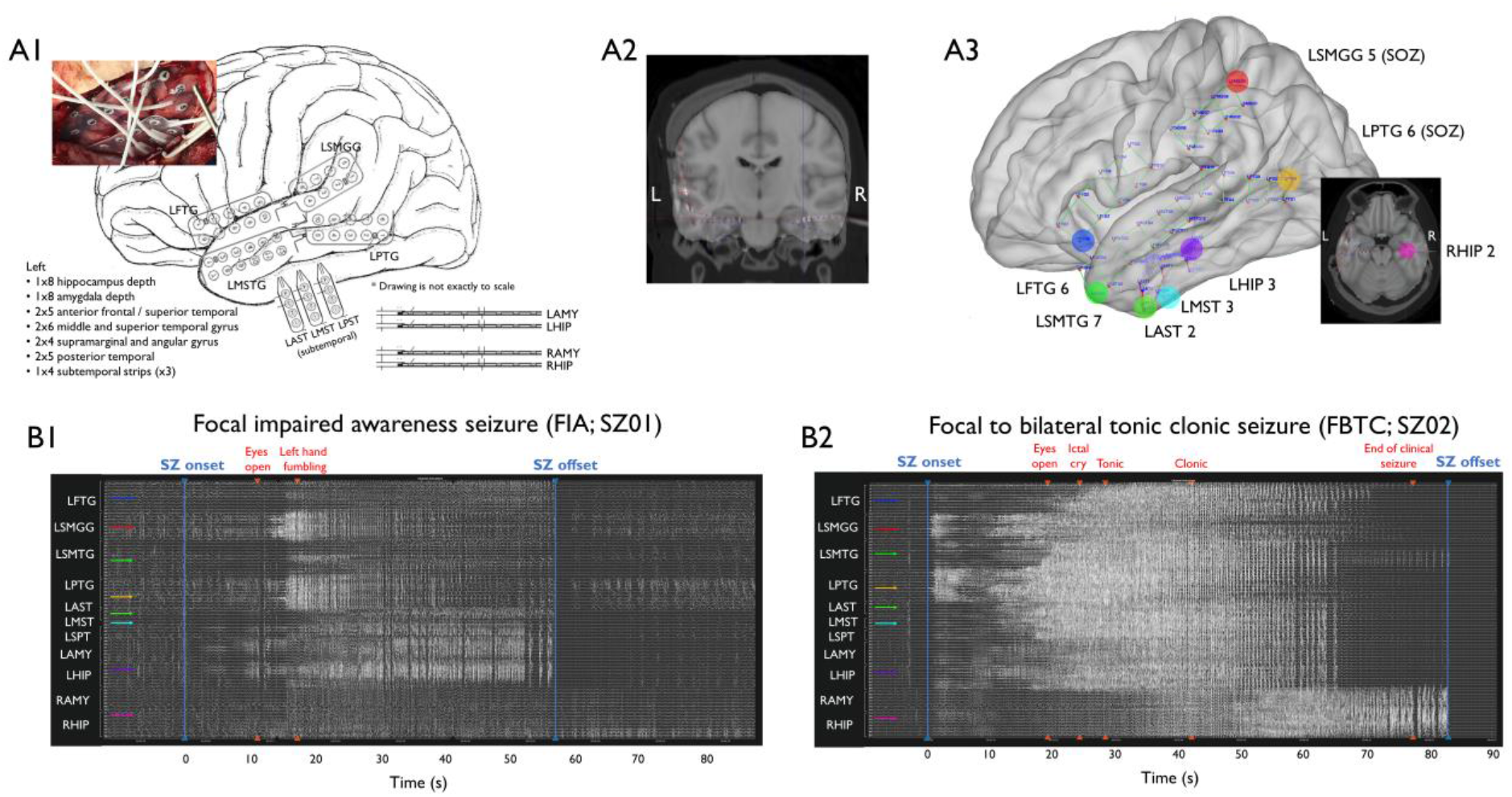
Exemplar 3D localization of intracranial electrodes (A) and iEEG signal for one FIA and one FBTC seizures (B) originating from a representative patient. (**A**) The surgical implantation and clinical electrode map (A1) are used to locate electrodes positions in the post-implantation CT and pre-implantation MRI images, aligned in MNI standard space using the iElectrodes software (A2); eventually, final electrodes localizations are displayed on a standard MRI template (A3). (**B**) Time courses of raw iEEG signals for a FIA (B1) and a FBTC (B2) seizure, along with marked behavioral events. Arrow colors indicate the colored electrodes displayed in A3. Contacts LSMGG 5 (in red) and LPTG 6 (in orange) were identified as belonging to the SOZ.

All iEEG data were acquired using either a subgaleal electrode (for SGDE) or a scalp electrode (for SEEG) as reference. This scalp electrode was placed in a fronto-central position (i.e. between Cz and Fz) for iEEG.org and EED, and to the right or left mastoid for UWHC. IEEG signal was preprocessed with customized scripts using routines from the EEGLAB toolbox version 14.1^39^ running on MATLAB 2016b (Mathworks Inc., Natick, MA, USA). A baseline consisting of minimum of 2 minutes (maximum 5 min) of pre-ictal iEEG signal was included (see Fig.1 B1-B2 for exemplar FIA and FBTC seizures). The raw signal was down-sampled to 400 Hz (including anti-aliasing filter) and band-pass filtered around 0.5 – 199 Hz using Hamming windowed sinc FIR filter. Line noise was removed when appropriate using the CleanLine algorithm (part of EEGLAB) on selected frequencies (mostly 60, 120 and 180 Hz). Channels and epochs with important artifacts (e.g. muscle activity) were rejected based on visual inspection.

### Power spectrum analyses and considered time periods

Power spectrum density was calculated for each contact using the ‘pwelch’ function with a 10 s window with 1 s overlapping window. Power spectrum values were then averaged over frequency bands of interest: SWA (1-4 Hz), beta (15-25 Hz) and high-gamma (HG; 80-150 Hz). In addition, B/D (B/D) was calculated as the quotient of beta power over SWA power.

SWA was used as a marker of sleep-like activities, as previously shown during FIA.^14^ Because B/D has been shown to differentiate sleep from wakefulness better than SWA within iEEG recordings,^40,41^ we considered B/D as an indicator of cortical activation. Considering that 80-150 Hz HG synchrony more specifically increases in actively seizing areas,^42,43^ we used this range of HG activity to quantify ictal activation.

Finally, we computed phase-locked high-gamma (PLHG; i.e. HG phased-locked to low 4-30 Hz oscillations) to obtain a marker of ictal activation validated to selectively increase in recruited areas.^18^ For each electrode, we obtained instantaneous phase and amplitude for high- and low-frequency components from the Hilbert transform of the respectively high-(80–150 Hz) and low-pass (4–30 Hz) filtered signals (windowed sinc FIR filter with Hamming window). The PLHG index was then computed within non-overlapping 1 s sliding windows as:

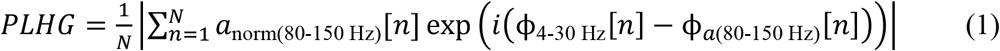

where *N* is the number of samples within the window, *a*_norm(80-150 Hz)_ is the instantaneous HG amplitude normalized by the average HG amplitude during baseline, *ϕ*_4-30 Hz_ is the phase of the low frequency component, and *ϕ*_a(80-150 Hz)_ is the instantaneous phase obtained from a second Hilbert transform applied to instantaneous HG amplitudes.^18,44^

To characterize the temporal evolution of different EEG features during seizures, the ictal period was split in two equal parts: the “first seizure half” corresponding to the seizure onset up to the midpoint of the seizure, and the “second seizure half” corresponding to the midpoint to the ictal offset. Note that a separate analysis described below looked at EEG correlates of behavioral generalization itself. Ictal values were normalized by pre-ictal baseline values for each seizure separately.

We performed group statistics using a linear mixed-effect (LME) model on normalized power values, with separate models fitting for each above-mentioned frequency band. Patient and seizure were entered as random effects, and seizure type, brain region and ictal period as fixed effects. The use of Restricted Maximum Likelihood (REML) estimates of the LME parameters allowed to account for the lack of independence between subjects and between seizures inherent to this dataset. The assumptions of the model were satisfied as the residuals showed Gaussian distribution. Statistical significance was evaluated at level *p*<0.05 and corrected for multiple comparisons using false discovery rate (FDR^45^). All analyses were performed in R (R Core Team, 2015), using the lme4 package^46^.

### Epileptogenicity index and SWA synchrony

To detect the spread of ictal recruitment, we quantified the number of channels crossing the energy ratio (ER) threshold between high and low frequencies used to compute the Epileptogenicity Index (EI).^28^ The EI is a validated measure of difference in timing of ictal recruitment between intracranial channels. It is typically used to determine the most likely SOZ by ranking channels according to the delay of their ictal involvement. We used ER threshold crossing to compute a proxy of the number of recruited channels during each seizure and compare the proportion of recruited channels across the whole brain and in each brain region between FIA vs. FBTC.

We also used timing information of ER threshold crossing to compare the (a)synchrony in ictal onset across iEEG channels during FIA vs. FBTC. Specifically, we defined the asynchrony parameter for a given seizure as the proportion of channels showing ictal recruitment (assessed with ER) more than 1 s apart from each other. To do so we computed for each seizure how many recruited channel units were more than 1 s apart then divided that number by the total number of channels (ratio in TableS3). As another marker of (a)synchrony, we quantified differences in the timing of peaks in SWA power (window size of 10 s and windows step of 1 s), and of high-gamma power (same parameters) for comparison. For each seizure, we assessed the proportion of channels that were more than 1 s apart regarding the timing of their SWA peak power. Finally, we also computed differences in timing of the peak amplitude of slow waves (SWs). SWs were detected within each channel using the procedure described by^47^ (1-4 Hz bandpass, third-order Chebyshev filter). We then assessed averaged SWs amplitude using sliding windows with window size of 10 s and windows step of 1 s and mark the position in time with maximal negative peak for each channel. For each seizure, we then calculated the proportion of channels presenting negative peaks that were at least 1 s apart from each other. We then computed group statistics for those measures of SWA asynchrony to compare FIA and FBTC (Table S3).

### Characterize the iEEG signatures of behavioral generalization

To characterize the electrographic changes accompanying generalization, we quantified iEEG brain activity changes preceding and coinciding with the onset of bilateral tonic phase. We selected a subset of 25 FBTC for which the onset of behavioral generalization (i.e. the start of the tonic phase) was known (from both UWHC and iEEG.org), and split the ictal period in pre- or post-generalization periods. To differentiate the markers of generalization vs. seizure onset, we contrasted SWA, B/D, HG and PLHG indices during the pre-ictal period, the pre-generalization period and the post-generalization periods.

To assess which iEEG indices were predictive of evolution of a seizure towards generalization, we compared SWA, B/D, HG and PLHG indices between the pre-generalization phase of FBTC and the first half of a subset of 57 FIA (coming from the same source as FBTC). FBTC pre-generalization period lasted on average 26±43s and FIA first half 39±51s (median±IQR), resulting in similar durations (*U*=837, *p*=0.2).

Although our analysis included exclusively iEEG signals from intracranial sources - thus limiting the contribution of muscle artifacts to high-frequency activity - we further wished to ensure that our results were not contaminated by extra-cerebral signals such as electromyogram activity (EMG).^48–50^ To investigate this point, we computed the contribution of EMG signals to iEEG channels filtered in HG band using linear regression in five FBTC recordings (including depth and surface iEEG channels from both left and right hemispheres) where EMG data was available (two from UW, three from EED). Because time-resolved behavioral data was not available in FBTC coming from EED, a proxy of generalization time was defined using timing of changes in PLHG. We first validated this measure in the 25 FBTC seizures for which behavior timing was available by calculating the average delay between the highest slope of increase in PLHG power and the behavioral generalization timepoint. On average, behavioral generalization occurred 12.87±2.11 s (mean±SEM) after PLHG highest increase slope (see Fig. 4D). To take into account global differences in amplitude, linear regression was preferred over bipolar contact subtraction; however bipolar montages between EMG and EEG led to similar results. We computed EEG/EMG synchrony for the whole ictal period and reported results separately for the pre-ictal, the pre-generalization and the post-generalization periods.

In order to examine the possible contribution of intra-vs extracranial sources and the possible presence of a subcortical third-driver, we also computed the contribution of deep vs. superficial iEEG signals in the HG band using linear regression. We included 17 FBTC for which both deep and superficial contacts were available (seven FBTC from UWHC, one FBTC from iEEG.org and nine FBTC from EED). In particular, we contrasted changes in synchrony in deep (amygdala/hippocampus) vs. superficial (neocortical) iEEG contacts between pre- and post-generalization periods. Deep and superficial iEEG contacts were normalized by their distance to avoid any bias on synchrony that could be attributed to different distance between contacts. The above-mentioned proxy for generalization time was applied for FBTC for which the point of behavioral generalization was unknown.

### Quantification of neuronal firing rates

To assess whether high frequency power changes at the macro-level in intracranial recordings of FBTC could indicate micro-level changes in neuronal firing, we used a unique dataset combining macro- and microelectrode recordings. We analyzed three additional FBTC seizures recorded in human epileptic subjects, where both iEEG and multi-unit activity were recorded, with Utah arrays located in areas remote from the SOZ (in the ictal penumbra, not recruited by the focal ictal process). We quantified local increases in firing rates compared to baseline, and their correspondence to sustained HG increases and behavioral generalization during FBTC.

Spike waveforms were detected on 30 kHz signal, filtered using a 1024^th^-order FIR bandpass filter from 300 to 5000 Hz, using a detection threshold of 4.5*σ*, where 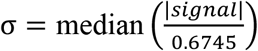 estimates the standard deviation of the background noise.^51^

From here, spike waveforms from the peri-ictal and ictal periods were examined separately, since traditional spike sorting methods have been shown to fail in ictal recordings.^52^ Waveforms from the peri-ictal period were sorted visually after *k-*means clustering using the UltraMegaSort2000 MATLAB toolbox,^53^ and noise artifacts were removed. Ictal unit firing was then analyzed using methods of Merricks *et al*.^54^. Spike waveforms from the ictal period were template-matched to sorted peri-ictal putative units if the principal component vector of the ictal spike fell within the convex hull of the peri-ictal unit’s spikes in principal component space. Further, a match confidence score from 0 to 1 was assigned to each matched ictal spike. For each peri-ictal unit, a Gaussian curve was fit to the distribution of voltages at each sampling point, then rescaled to a maximum height of one. Then, for each ictal spike matched to that unit, the spike’s voltages were mapped to a point on each Gaussian, and the average of these values over each sampling point resulted in the match confidence score.

We used probabilistic multi-unit firing rates based on putative neurons recorded by Utah microelectrode array^54^. The instantaneous probabilistic firing rate was calculated by convolving a discrete Gaussian kernel of standard deviation of 200 ms with the detected spike train, where each detection is scaled to its match confidence. Therefore, the probabilistic firing rate could not be artificially increased by an increase in background noise or by a single spike waveform matching with multiple putative units. The average probabilistic firing rate over a period of time was calculated by dividing the sum of all match confidence scores for detected units in that period by the length of the period. This average firing rate was used to quantify multi-unit activity in the pre-ictal (20 s. before onset to seizure onset), pre-generalization (seizure onset to behavioral generalization), and post-generalization (behavioral generalization to seizure offset) periods. In addition, for each putative single unit, spike times of waveforms with >50% match confidence were displayed across the seizure epoch in a raster plot. Units were ranked for display based on overall firing rate during the seizure epoch. Code for firing rate calculation and raster plots can be found at https://github.com/edmerix/NeuroClass.

Additionally, to compare local neuronal firing in the Utah array region with global dynamics during seizure, the PLHG metric was computed on both macro- and microelectrodes for these subjects as above. After removing faulty electrodes and those near lobar/SOZ boundaries, macroelectrodes were partitioned by region into frontal, temporal, parietal, and SOZ groups, along with a single electrode nearest the Utah array. Mean PLHG values over the seizure epoch were then calculated for each electrode group, including the group of Utah array microelectrodes.

### Data availability

Data supporting the findings of this study are available from the corresponding author upon reasonable request. Data from iEEG.org and EED are available online.

## Results

### Markers of LOC distinguish FBTC from FIA

Behavioral assessment of 42 FIA where video was available (from UWHC and Mayo Clinic) revealed verbal unresponsiveness in 80% of tested seizures, motor unresponsiveness in 79% and amnesia of seizure events in 84%. In 46% of tested FIA, patients failed to interact with examiner in any way (CSS 3) and in 79% they were not aware of having a seizure during the event (CSS 4).

Splitting LOC scoring in two equal halves in FIA (30±18 s) revealed signs of a progression across time. Specifically, while 73% patients were not aware of having a seizure in the first half of FIA, this significantly increased to 97% in the second half (Fisher’s exact test: *P*=0.007). Verbal unresponsiveness, motor unresponsiveness and interaction with examiner also seemed to become more impaired in the second half (from 68% to 70%, from 54% to 74% and from 45% to 50% respectively), but these variables did not reach statistical significance.

In the 22 FBTC for which the behavior before generalization could be scored (duration of pre-generalization phase: 58±67 s), pre-generalization behavior was characterized by verbal unresponsiveness in 89% and motor unresponsiveness in 93%. Interaction with examiner was impaired in 90% (CSS 3) and patients were not aware of having a seizure (CSS 4) in 91% of seizures. Contrasting these outcomes with the first half of FIA, we found significantly more patients with motor unresponsiveness (*P*=0.01; Fisher) and impaired interaction with examiner (*P*=0.001; Fisher). Post-generalization, all FBTC patients were unresponsive and unable to sustain any interaction, which was the case in respectively 74% (*P*=0.008; Fisher) and 50% (*P*<0.001; Fisher) of patients in the second half of FIA.

### FBTC are accompanied by more widespread increases in sleep-like activities and in cortical activation

The linear mixed-effects model for SWA revealed no initial change from baseline during the first half of FIA (Fig. 2 left panel; see Table S1 for all *Z* and *P* values). A significant SWA power increase was subsequently seen in all brain areas during the second half (e.g. increase from 1.01±0.03 to 1.67±0.03 in SOZ; *Z*=-22.35, *P*<0.001). In contrast, in FBTC, a significant SWA power increase was already seen in SOZ, temporal and frontal regions during the first half (*P*<0.001; see Table S1 for all *Z* and *p* values). SWA further increased during the second half of FBTC in all brain areas (e.g. increase from 1.30±0.05 to 2.77±0.06 in SOZ; *Z*=-32.5, *P*<0.001). While between-seizure contrasts were not significant for the first seizure half, significantly more SWA in parieto-occipital regions was seen during the second half of FBTC compared to the second half of FIA (*P*=0.001).

**Fig. 2.**
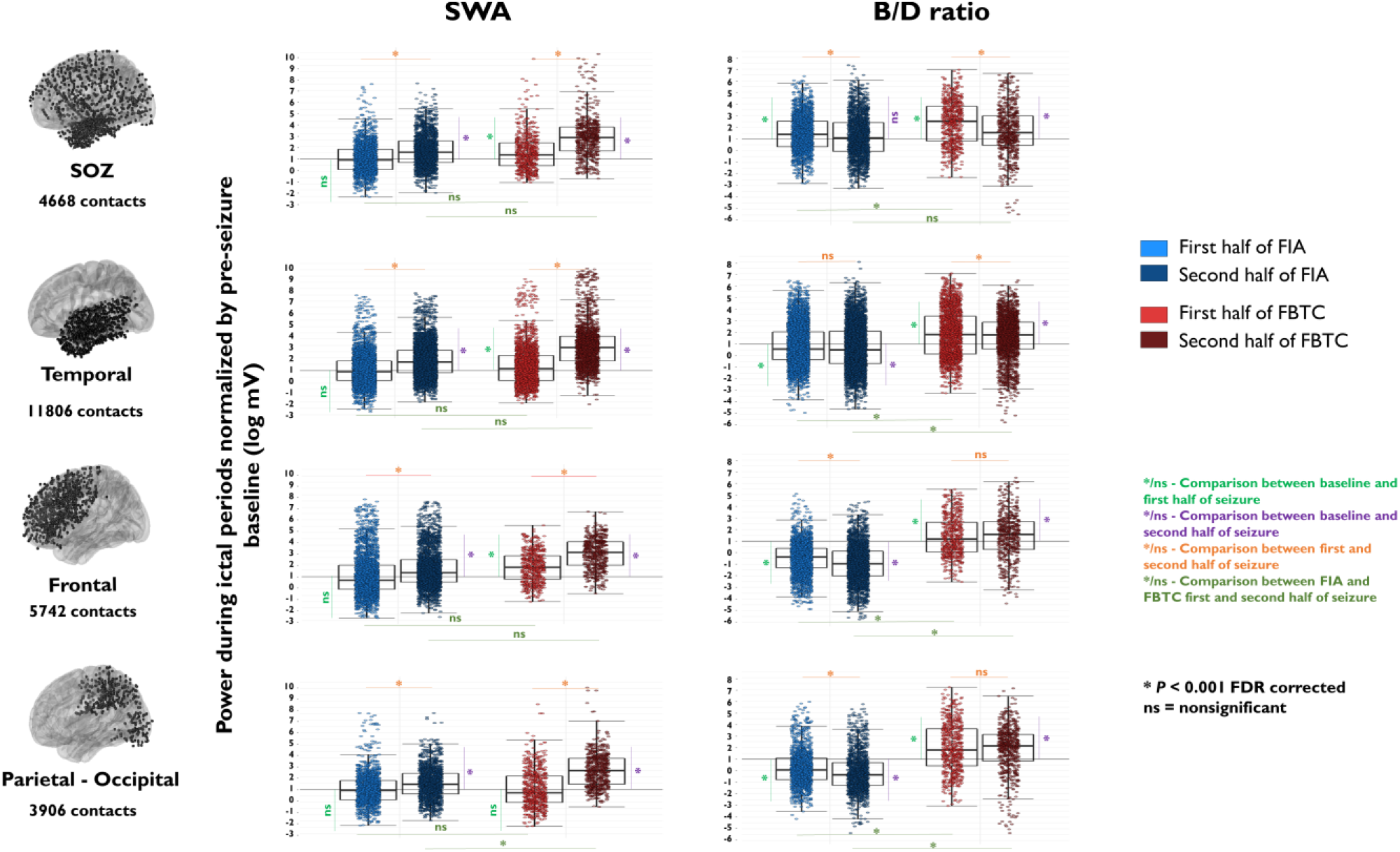
Group results for SWA power and B/D during both FIA and FBTC, split by brain region and ictal period. Each dot represents the log of normalized power value (normalized by baseline activity) for an electrode contact. FIA are displayed in blue and FBTC in red, with lighter colors indicating the first ictal period, and darker colors indicating the second ictal period. Black horizontal lines indicate values of pre-ictal baseline activity. These results suggest that SWA increases and B/D decreases are prominent during the second ictal period during FIA. In contrast, both SWA and B/D increase in the whole brain starting at the onset of FBTC.

During the first half of FIA, B/D decreased in all brain regions compared to baseline except in SOZ, where it was increased (Fig. 2 right panel; see Table S1 for all *Z* and *P* values). A significant B/D decrease was further seen between first and second half of FIA in frontal and parieto-occipital regions (−0.16±0.05 in parieto-occipital areas, *P*<0.001; −0.93±0.03 in frontal areas, *P*<0.001). In sharp contrast, B/D showed widespread and consistent increases during FBTC. Indeed, during the first half of FBTC, B/D increased in all brain regions, and most prominently in SOZ (e.g. 2.37±0.07 in SOZ, *P*<0.001; 1.84±0.05 in temporal areas, *P*<0.001). B/D then remained elevated in frontal and parieto-occipital regions, while it decreased in SOZ and temporal areas. Between-seizure contrasts showed increased B/D for FBTC compared to FIA during both halves of the seizures and in all brain regions.

### FBTC are accompanied by higher HG activity and cross-frequency coupling in many cortical areas

During FIA, HG power initially increased in SOZ, parieto-occipital and temporal regions, while it decreased in frontal areas (Fig. 3 left panel; see Table S2). HG power remained increased in SOZ, temporal and parieto-occipital areas during the second half of FIA, while it returned to baseline in frontal areas. In contrast, during the first half of FBTC, HG power showed more than two-fold increase in all brain regions as compared to baseline, which then further increased more than three-fold from baseline in all areas in the second seizure half (*P*<0.001; Table S2). Between-seizure contrasts revealed significantly higher HG power in FBTC compared to FIA during both seizure halves and in all brain regions.

**Fig. 3.**
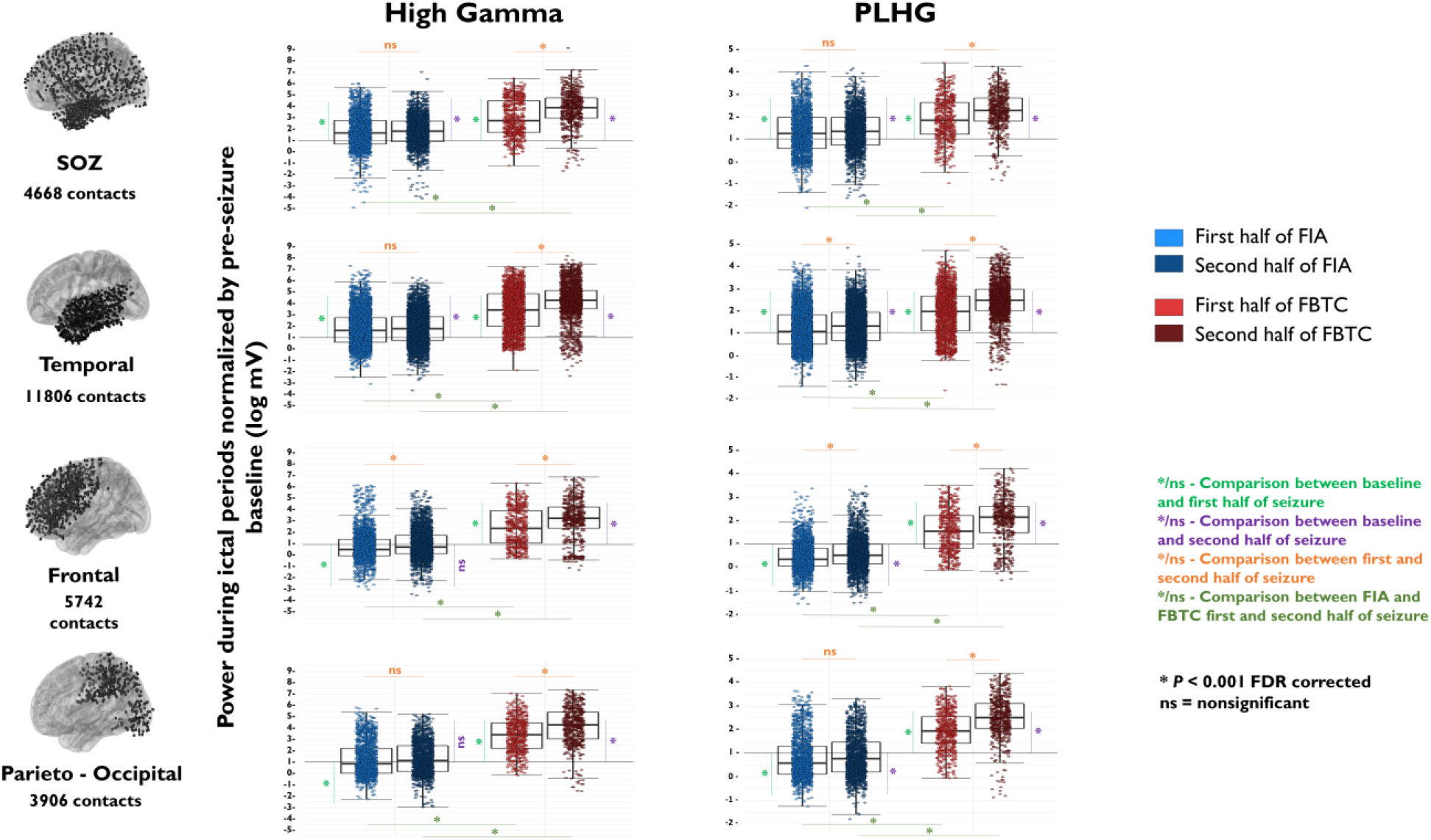
Group results for High Gamma (HG) power and Phase Locked High Gamma (PLHG) during both FIA and FBTC, split by brain region (SOZ, temporal, frontal and parieto-occipital) and ictal period (first and second half of seizures). Each dot represents the log of normalized power value (normalized by baseline activity) for an electrode contact. FIA are displayed in blue and FBTC in red, with lighter colors indicating the first ictal period, and darker colors indicating the second ictal period. Black horizontal lines indicate values of pre-ictal baseline activity. These results suggest that HG and PLHG increase in SOZ and temporal lobe but decreases in the rest of the brain during FIA. In contrast, both HG and PLHG diffusely increase starting at the onset of FBTC, and further build up as FBTC progress. HG and PLHG were significantly higher during FBTC than during FIA for all brain areas and all ictal periods.

During the first half of FIA, PLHG increased from baseline in SOZ and temporal areas (*P*<0.001; Fig. 3 right panel; Table S2) while it decreased in frontal and parieto-occipital areas (*P*<0.001). During the second half, PLHG continued to show a mild increase compared to baseline in temporal areas, remained decreased in frontal and parieto-occipital areas, while was more variable in the SOZ (1.35±0.02, p=0.195 for SOZ, 1.30±0.01, *P*<0.001 for temporal, 0.60±0.01, *P*<0.001 for frontal, and 0.86±0.02, *P*=0.024 for parieto-occipital). In contrast, during the first half of FBTC, PLHG increased by 1.5 times from baseline for all brain areas (Table S2b). During the second half, PLHG was further increased (*P*<0.001 for all brain areas when comparing to baseline values, e.g. 2.52 fold increase in the parieto-occipital area). Between-seizure contrasts revealed significantly higher PLHG in FBTC compared to FIA for both seizure halves in all brain areas (Table S2).

### Widespread but asynchronous ictal recruitment during FBTC

The energy ratio (ER) analysis also showed that more channels were recruited in the ictal process during FBTC compared to FIA (69±4% and 45±6% respectively, *P*<0.001; Table S3). Overall, channels that least frequently passed ER during FIA were located in the parietal lobe (29% vs 39-60% in other lobes, *P*<0.001) and during FBTC, in the limbic network (58% vs 62-78% in other lobes, *P*<0.01).

The analysis of the timing of ER threshold crossing revealed that the recruitment of channels into the ictal process was asynchronous both during FIA (example in Fig.4A1) and FBTC (example in Fig.4A2). Interestingly, we found that while more channels were recruited during FBTC, significantly more clusters of channels were also recruited more asynchronously (with a higher proportion of channels crossing ER threshold more than 1s apart: 23±13% in FBTC vs. 10±7% in FIA, *P*<0.001, Table S3).

**Fig. 4.**
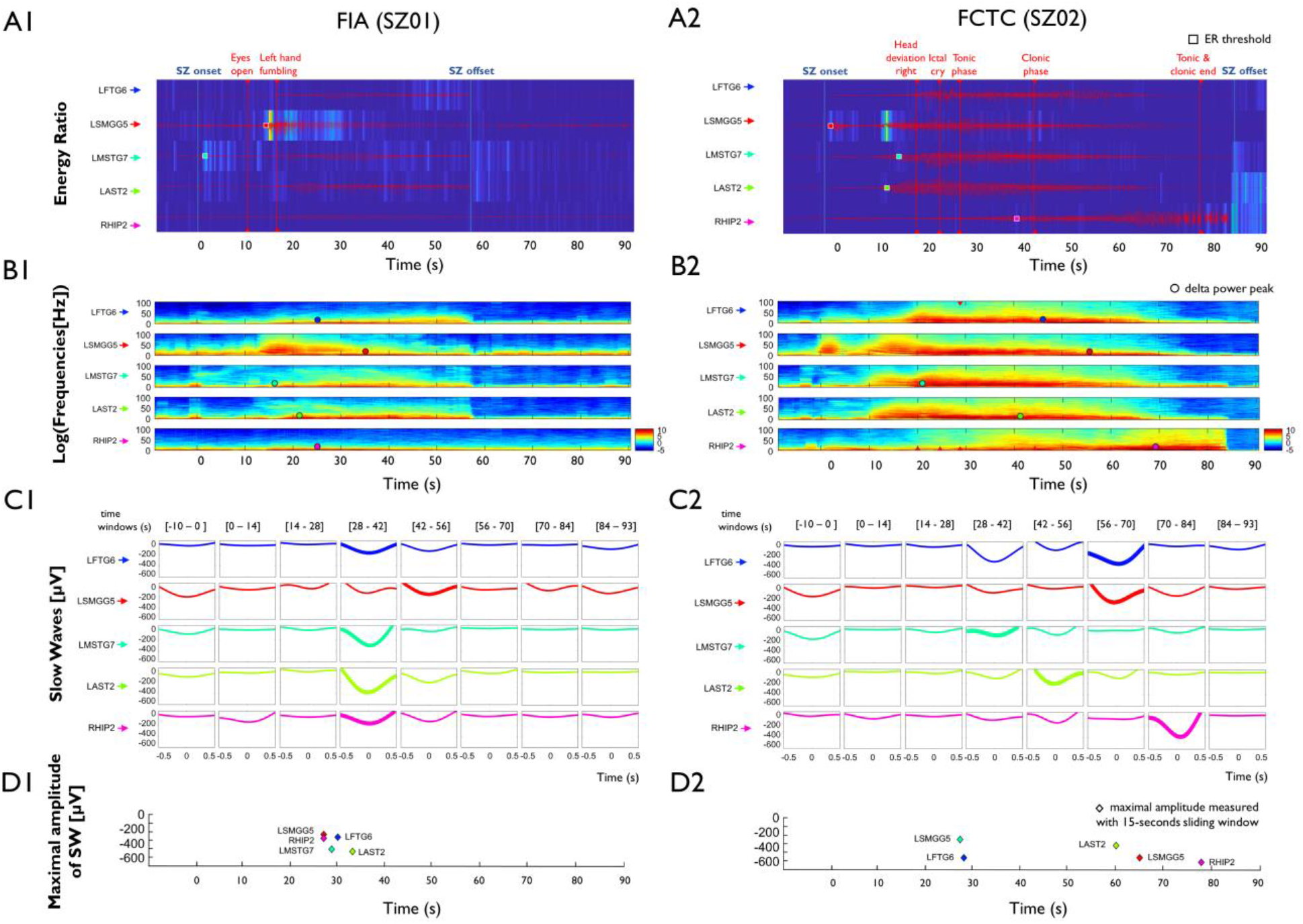
Asynchrony of channel recruitment, SWA power and slow wave (SW) amplitude peaks during FIA vs FBTC. Five representative channels are displayed for the same representative FIA (left panel) and the FBTC (right panel) used in previous figures. Panel A displays the time points where the Energy Ratio (ER) threshold was exceeded as dots with different colors corresponding to each channel. Channels were recruited asynchronously in both cases, but with less channels recruited during FIA than during FBTC. At the time of the behavioral generalization (red line), only a partial recruitment could be observed. Panel B displays the time-frequency representation for the same seizures, with asynchronous peaks in SWA power marked with dots of colors corresponding to each channel (see Fig. S1 for the timing of high gamma power peaks). Panel C displays average slow waves (SWs) detected at specific time intervals, with windows with maximal SW amplitude highlighted in BOLD. Panel D displays the timing of negative SW amplitude peaks with dots in colors corresponding to the same channels than in other panels. Note that the timing of occurrence of SW amplitude peaks is especially asynchronous during FBTC.

To compare differences in synchrony of ictal patterns during FIA vs FBTC, we examined the dynamics of time-frequency activity across channels (Fig. 4B). During both seizure types, we observed asynchronous increases in SWA power across different channels, with most channels presenting SWA power peak during the second half of the seizure. SWA power peaks were more asynchronous during FBTC than during FIA (34±16% and 28±16%, *P*=0.025, Table S3). Similar to SWA, the timing of the most negative amplitude of individually detected slow waves (SW) most often occurred during the second half of the seizures (mean time of 75±12 s for FBTC, and 63±17 for FIA). The timing of occurrence of most negative SW amplitude was again more asynchronous during FBTC as compared to FIA (42±18% vs. 35±16% of SW amplitude peaks crossing threshold more than 1 sec apart, *P*=0.012, Table S3).

In contrast, during both seizure types we observed synchrony of high-gamma power peaks across channels (Fig. S4), which was specifically prominent during FBTC. Interestingly, during FBTC HG power also increased across the ictal period in channels that were not recruited by the seizures (did not pass the ER threshold; Tables S11 and Fig. S4).

### Extra-temporal beta/delta ratio and PLHG increases distinguish FBTC pre-generalization from the first half of FIA

Comparing FBTC pre-ictal period (baseline) to FBTC pre-generalization period revealed a significant SWA power decrease in SOZ and temporal areas (*P*<0.001; see Table S7 and Fig. S2). Additionally, B/D increased in SOZ and decreased in parieto-occipital and temporal areas (*P*<0.001 for SOZ and parieto-occipital areas, p=0.023 for temporal areas; see Table S7 and Fig. S2). SOZ and temporal areas also showed a significant increase in HG power in the pre-generalization period (*P*<0.001), while parieto-occipital and frontal areas remained at baseline level (Table S8a). Finally, PLHG showed significant increase in SOZ and decrease in parieto-occipital and frontal areas (see Table S8b).

In comparison to the first half of FIA, B/D was significantly increased during pre-generalization in FBTC in SOZ, frontal and parieto-occipital areas (*P*<0.01; Table S9b). Interestingly, PLHG was also significantly higher in FBTC pre-generalization as compared to FIA first half, both in SOZ and in parieto-occipital areas (*P*<0.001 and *P*=0.007 respectively; Table S10b). In contrast, there was no difference between the early phase of both seizure types in any brain area for SWA (Table S9a) or HG (Table S10a).

### Extra-temporal B/D and PLHG increases distinguish FBTC (pre-generalization) and FIA (first half)

In comparison to the first half of FIA, B/D was significantly increased pre-generalization in FBTC in SOZ, frontal and parieto-occipital areas (*P*<0.01; Table S9b). Interestingly, PLHG was also significantly higher in FBTC pre-generalization as compared to FIA first half in SOZ and in parieto-occipital areas (*P*<0.001 and *P*=0.007 respectively; Table S10b).

In contrast, there was no difference between the early phase of both seizure types in any brain area for SWA (Table S9a) or HG (Table S10a).

### A whole-brain increase in high-gamma activity coincides with behavioral generalization

The above-mentioned group analysis performed on all seizures suggested a widespread increase in HG power and PLHG during FBTC compared to FIA, which built up from the first to the second half of the seizures. To further characterize if such a late build-up in high-frequency activity was related to the occurrence of behavioral generalization itself, we examined the correspondence between spectral power time courses and ictal behavior. During FIA, a minimal increase in HG power could be seen in the SOZ, peaking in the middle of the seizure (example in Fig. 5, right panel). During FBTC, stronger increases in HG power and PLHG were consistently seen in the SOZ from the seizure onset (example Fig. 5, left panel), which further invaded all channels, with the sharpest slope for a whole-brain increase closely matching the time of behavioral generalization onset (see example in Fig. 5B). Proof-of-principle analyses of the 25 FBTC with behavioral scoring showed that there was a significantly higher time concordance between the point with maximum slope of HG increase and behavioral generalization (21±3.7 s from behavioral generalization; Table S8c) compared to seizure onset or offset. Interestingly, the temporal concordance with behavioral generalization was higher with PLHG (12.9±8 s) than for SWA (37±40 s; Table S6c).

**Fig. 5.**
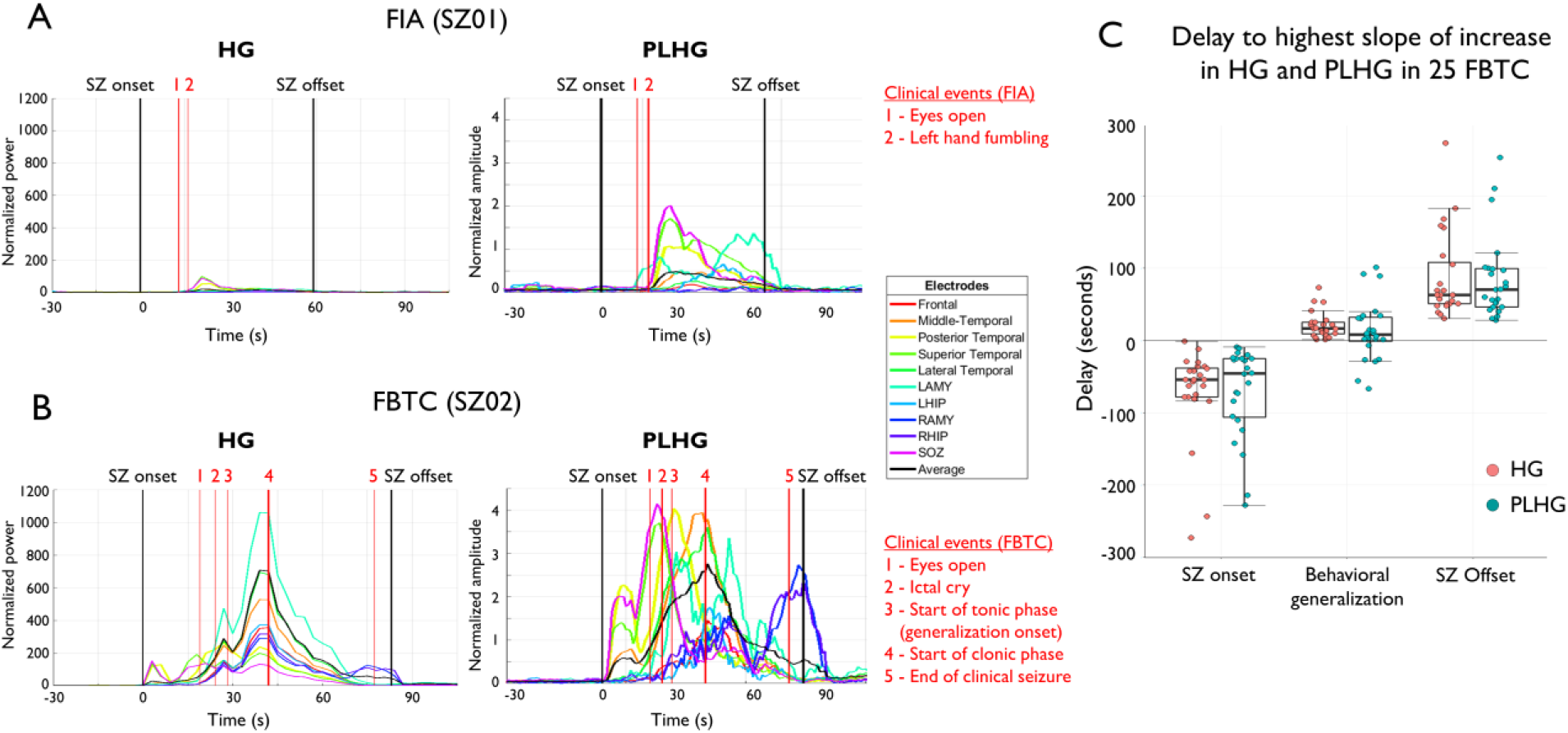
Temporal evolution of high frequency rhythms (HG and PLHG) in two exemplar FBTC and FIA from the same patient (same as in Fig. 1), displayed along relevant behavioral events. In FBTC (panel B), the widespread increase in HG and PLHG peaking at generalization point (event 3) can be clearly seen. Panel D displays group results (from 25 FBTC) for the average delay between seizure start, generalization point and seizure end to the highest slope increase in HG power and PLHG.

### Increase in HG is not a muscular artifact

A linear regression between EMG and iEEG channels revealed that EMG accounted for only on average 1.31%±0.34 % (range 0.01-12%) of explained variance in the iEEG across the whole ictal period (Fig. 6A; Tables S6). In contrast, iEEG channels on average showed a shared variance of 43±4 % (range 12-91%). In fact, iEEG channels shared significantly higher variance after generalization than before generalization or during baseline compared to iEEG vs EMG (*t*_*(19)*_=2.28, *P*=0.351 for baseline; *t*_*(19)*_=9.4, *P*<0.001 for pre-generalization; *t*_*(19)*_=10.71, *P*<0.001 for post-generalization). This suggests a minimal or non-existent contribution of muscle activity. Linear regression between deep vs. superficial contacts revealed higher synchrony between deep contacts (amygdala, hippocampus) than between superficial contacts, especially during the post-generalization period (47±33% then 61±23% for superficial contacts, 82±35 % then 81±20% for deep contacts; Fig. 6B, Table S6).

**Fig. 6:**
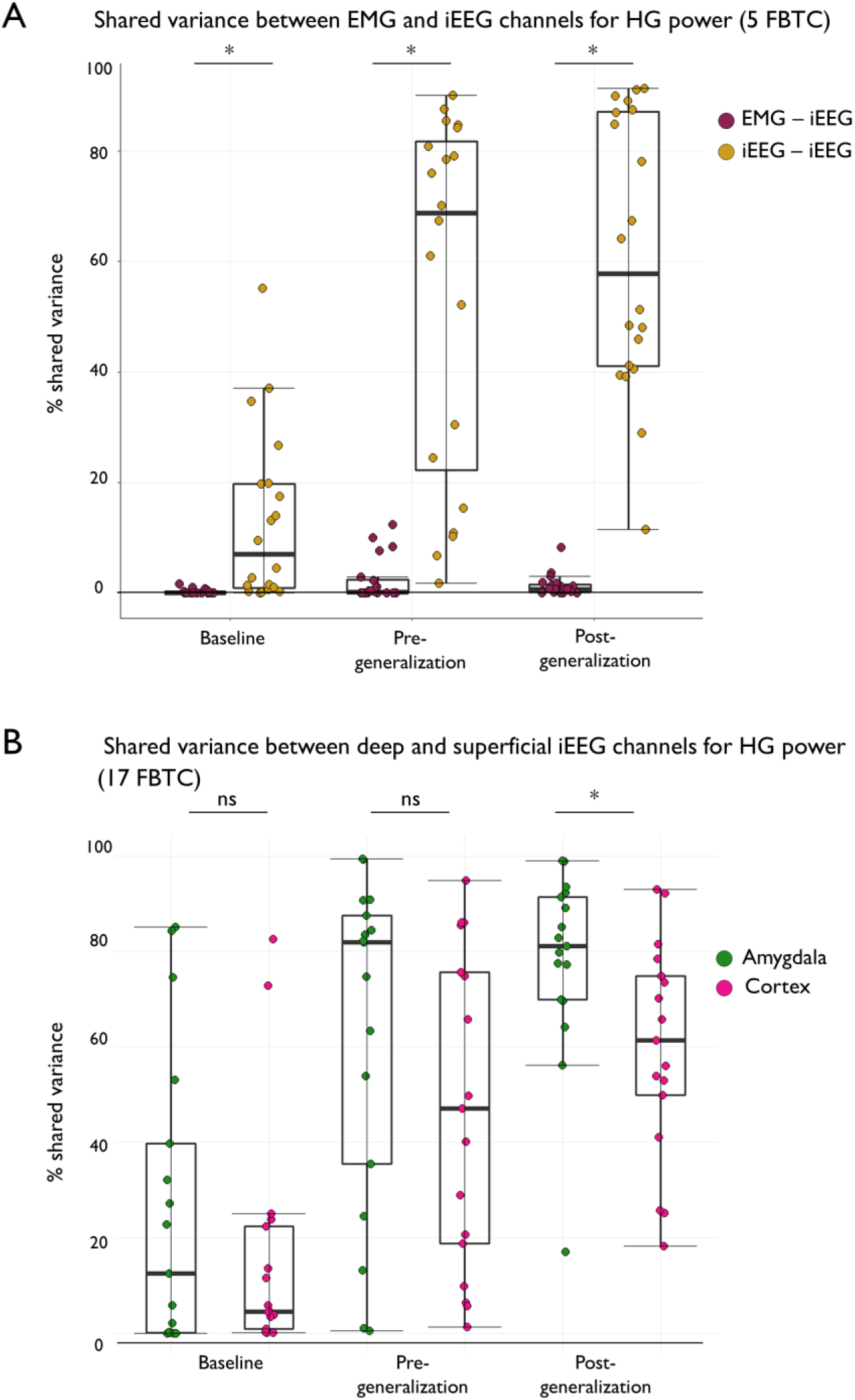
Shared variance in HG power between EMG and iEEG electrodes, and between deep versus superficial iEEG electrodes during FBTC. (A) The portion of HG signal in iEEG electrodes that is explained by EMG activity is negligible compared to the variance explained by other iEEG electrodes across all FBTC periods (baseline, pre-generalization and post-generalization). (B) Deep electrodes – especially in the amygdala and hippocampus – share more common variance in their HG signals than superficial cortical electrodes during the post-generalization period. This finding suggests a potential deep source as HG power generator during this period.

### Increased neuronal firing rates accompany HG increases in non-SOZ areas during FBTC

To obtain a direct demonstration of the neuronal basis for high-frequency signal changes during FBTC, we used simultaneous iEEG and single-unit recordings in three FBTC recorded with Utah microelectrode arrays in areas located in the ictal penumbra (>2 cm from the SOZ) (see Fig. 7). Before generalization time, probabilistic multi-unit firing rate changes compared to baseline were variable: the average increased in one FBTC and decreased in two others (+51.5%, −43.4% and −48.5% change, respectively). In contrast, probabilistic multi-unit firing rates showed a sustained and consistent increase after the point of behavioral generalization compared to baseline (+215.8%, +83.4%, and +234.6% change, respectively). Increases in firing rates at generalization onset were accompanied by increases in HG within the whole brain (as observed in the other FBTC studied).

**Fig. 7.**
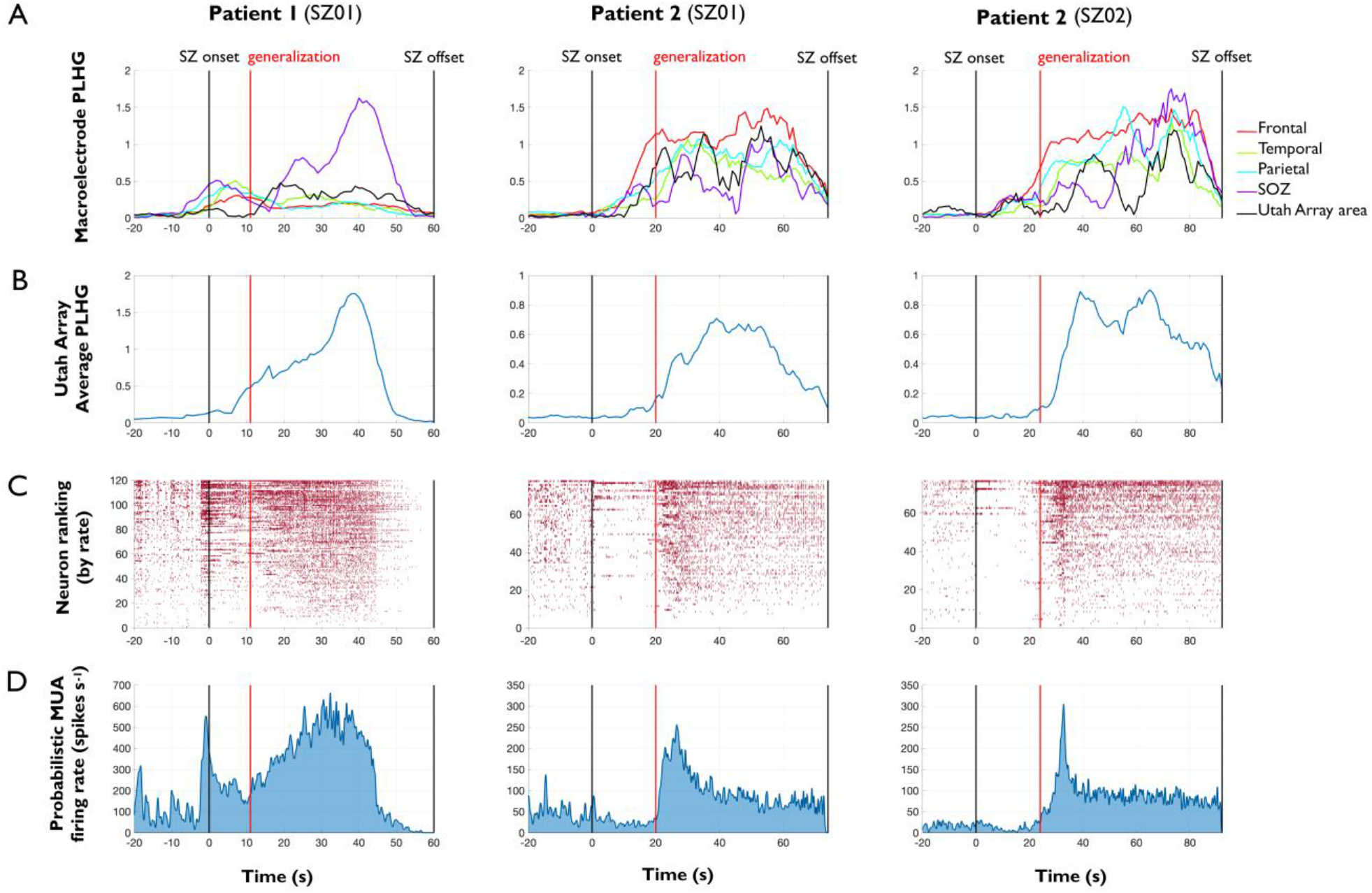
Temporal evolution of PLHG and neuronal firing during three FBTC from two patients implanted with Utah microelectrode arrays in areas remote from the SOZ. (A) PLHG for macroelectrodes in various locations across the brain. PLHG values in the SOZ and near the Utah array come from single distinct contacts, while PLHG values from frontal, temporal, and parietal areas were averaged over several contacts in the corresponding lobe (see Fig. S3). (B) Average PLHG calculated across all good Utah array channels. (C) Raster plot of single unit firing times ordered by firing rate. Only spikes with match confidence of 50% or higher are plotted. (D) Probabilistic firing rate for the population were calculated over the seizure epoch. These results show similar high frequency increase at the macro level - with increased PLHG - and at the micro level - with increased neuronal firing - and this both for ictal onset and behavioral generalization.

These unit firing rate increases occur alongside PLHG increases both in the iEEG channels close to the Utah array and in Utah array micro-electrode channels (see Fig. 7). MUA and PLHG appeared linked - except during brief peaks of multi-unit firing that do not show corresponding peaks of PLHG. This is sensible, as PLHG is predicted to especially increase due to highly synchronous firing, but not all firing.^42^ Therefore, PLHG may not track multi-unit firing when firing is asynchronous, or when highly synchronous firing increases in frequency without increasing synchrony. Nevertheless, the tendency of these values to match over sustained periods supports the hypothesis that widespread PLHG increases are neuronal in origin.

## Discussion

We found that unlike during FIA, the temporal evolution of seizures during FBTC is accompanied by a diffuse increase in markers of cortical activation (B/D, HG, PLHG) and ictal recruitment (number of channels crossing ictal ER threshold). Specifically, a whole-brain increase in HG power accompanied behavioral generalization onset, which was most synchronous in deep iEEG channels, could not be accounted for by EMG, and was accompanied by increased multi-unit firing rates in areas remote from SOZ. Overall, these findings suggest different mechanisms for LOC during FBTC compared to FIA, with an increase in cortical activation and ictal recruitment rather than sleep-like activities. Interestingly, the maximum of synchrony in deep iEEG sources at the time of generalization (especially strong in amygdala) suggests a potential contribution of a subcortical source.

Both FBTC and FIA were associated with increased SWA during the second half of the seizures, when consciousness is most impaired.^10,55^ During the second half of the seizures, SWA power was found to be higher in FBTC than in FIA in parieto-occipital cortex. This finding is in line with recent studies on the neural correlates of sleep dreams^56^ and with clinical and neuroimaging evidence^57^ suggesting an important role of parieto-occipital cortex in human consciousness.

Our behavioral results revealed a severe LOC during FBTC seizures, and a milder alteration in FIA. It is worth noting that while during a majority of FIA, patients were not aware of having a seizure and were not responding to verbal and motor commands, they could often still interact with the examiner in a minimal way. Interestingly, even in the pre-generalization period of FBTC, responsiveness was more strongly impaired. These results confirm previous reports showing a moderate consciousness impairment in FIA and a more complete one in FBTC.^10–12,33,58^

Our results confirm and extend in a larger dataset the previous observation of widespread increases in cortical SWA during FIA of temporal lobe onset.^14^ During FIA, increased SWA in frontal and parieto-occipital areas was indeed sleep-like: it was accompanied with a decrease in B/D – which reliably differentiates physiological sleep from wakefulness in iEEG recordings.^40,41^ During FBTC, in contrast, B/D increased within the whole brain. The finding of widespread cortical activation fits with previous studies using electroconvulsive therapy in humans and with animal models showing diffusely increased brain metabolism and fMRI BOLD signal during FBTC.^14,21,59^ The fact that some individual channels showed B/D increases before they were actively recruited into ictal rhythms, and that this phenomenon occurs specifically in FBTC and not in FIA suggest that increased beta-delta ratio may at least in part be related to the activation of a third-driver source, potentially of subcortical origin. Furthermore, our synchrony analyses revealed that while SWA power developed asynchronously throughout seizures, HG was very synchronous, suggesting once again a possible subcortical main driver for whole-brain increases in high frequency activity during FBTC.

We also found a more widespread ictal activation during FBTC than during FIA. Indeed, FIA displayed PLHG increases which were mostly restricted to SOZ, while PLHG decreased in extra-SOZ brain regions. In contrast, PLHG increased early and diffusely in the whole brain during FBTC and further built up with FBTC progression. Because PLHG increases reliably indicate areas that are recruited into ictal firing,^24,25^ its progressive evolution during FBTC provides strong support for a more widespread ictal involvement than during FIA. Additionally, we found significantly more channels passing a validated ER threshold for ictal involvement during FBTC than during FIA. The higher cortical recruitment during FBTC evidenced here using two independent quantitative markers of ictal recruitment is in line with previous studies in smaller samples using less specific markers such as the visual detection of HFOs.^19^ This finding may explain longer-lasting cognitive consequences of FBTC and through the induction of plastic changes, its association with poorer surgical outcomes.

Interestingly, while more channels were recruited into the ictal process during FBTC than during FIA, we also found more asynchrony between clusters of channels. This was found using both the quantification of each channel’s ictal onset by crossing of the ER threshold,^28^ and the inspection of later ictal dynamics for SWA and SW amplitude peaks. This observation questions the fact that LOC during FBTC may be related to an increase in synchrony within cortical signals, as suggested in.^60^ It also points to possible association between the occurrence of FBTC and the development of multiple intracranial epileptic foci.^61^ The tools developed in the present work may be used in future studies to assess if the number of independent foci recruited during either FBTC and FIA differentially predicts multifocal seizures and poor surgical outcomes in epileptic individuals.

The electrophysiological hallmark of behavioral generalization was a widespread increase in HG power. This HG increase at generalization onset was especially synchronous in deeper channels (amygdala, and to a smaller extent, hippocampus). This suggests that deep sources particularly well-connected to limbic areas – such as arousal centers in the brainstem or basal forebrain – may be involved in spreading cortical activation during the generalization process. Unilateral blockade of inhibitory GABA neurotransmission in the basal forebrain is able to trigger bilateral limbic motor seizures in the rat.^62^ The involvement of subcortical structures in seizure generalization is also supported by a previous SPECT study demonstrating increased cerebral blood flow in the brainstem and basal ganglia during FBTC compared to FIA.^59^ This hypothesis is also in line with early work in cats suggesting that electrical activation of the brainstem can rapidly induce widespread increases in markers of cortical activation^63^ while its ictal involvement can generate tonic posturing^64^ and bilateral convulsions.^65^ Another potential candidate for the subcortical mediation of seizure generalization might be the zona incerta. Indeed, rodent studies showed that high intensity cholinergic stimulation of the zona incerta leads to generalized seizures with highest probability amongst all other subcortical sites.^66^ The zona incerta^67^ is a central relay of communication between the thalamus and the brainstem and presents especially rich interconnections with bilateral intralaminar and higher order nuclei of the thalamus.^67,68^

In the five patients where EMG channels were available, we found no meaningful contribution of EMG to HG signals, comparing EMG channel to deep and superficial channels from left as well as right hemispheres (average shared variance <5%). This finding suggests that as the HG increase that is observed during FBTC postictal states,^48,49^ the iEEG HG activity increases observed during and after generalization phase cannot be accounted by increased EMG activity. Findings of higher synchrony in deep iEEG contacts, further away from the scalp, and of only partial synchrony between cortical iEEG channels during the post-generalization phase also plead against an artifact as the primary source for the observed increases in HG signal.

To further ascertain of the neuronal origin of HG power increases, we quantified MUA data during three human FBTC. We found that increases in HG during FBTC were indeed accompanied by sustained increases in neuronal firing even in areas remote from the SOZ. Taken together, these findings suggest that during FBTC – unlike LOC during FIA – is accompanied by widespread increases in neuronal activation throughout the brain. Of note, frequent FBTC recruiting a large number of areas may favor Hebbian plastic changes and secondary potentiation of multiple areas in the cortex, further favoring conditions for secondary epileptic foci to emerge, even in non-recruited brain areas. Such changes could explain worsened surgical outcomes and global cognitive impairment found in patients with frequent FBTC seizures.

This study has several important limitations. Only nine patients had both FBTC and FIA seizures. However, electrode coverage and demographics were similar between patients with FIA and FBTC at the group level, with broad electrode coverage and a large number of included seizures in both cases. Only three FBTC were recorded with multi-unit activity recordings, and the present findings should be confirmed in additional datasets with recordings within and outside the SOZ. Finally, the evidence we have for subcortical third driver(s) is only indirect; animal studies may be more suited to test the contribution of various subcortical structures to seizure generalization and to explore various neuromodulation strategies. In combination with brain activity recordings, future studies should aim at probing behavior continuously throughout seizures. However, continuous behavioral sampling is extremely difficult to perform during FBTC. Retrospective collection of phenomenal experiences remembered by patient may provide useful complementary information about LOC in patients who are behaviorally unresponsive.^12^

## Conclusion

In summary, our results show that FBTC are characterized by widespread increases in high-frequency activity and neuronal firing, which further progressed after onset of behavioral generalization. This high-frequency activity was most synchronous in deep iEEG channels, hinting to the possible contribution of sub-cortical drivers. These findings suggest that LOC during human FBTC may occur through a different mechanism than during FIA, with the presence of widespread increase of neural activation throughout the cortex.

## Abbreviations

B/D: B/D
FBTC: Focal to Bilateral Tonic Clonic
FIA: Focal Impaired Awareness
HG: High Gamma
iEEG: intracranial EEG
PLHG: Phased-Locked High Gamma
SWA: Slow Wave Activity

## Acknowledgements

The authors would like to thank Pei-Ning Peggy Hsu for help with clinical information and William Marshall for statistical advice.

