## Supplementary material for "Distinct signatures of loss of consciousness during Focal Impaired Awareness versus Focal to Bilateral Tonic Clonic seizures": SupplMat_figures

### SUPPLEMENTARY MATERIALS

#### Figures

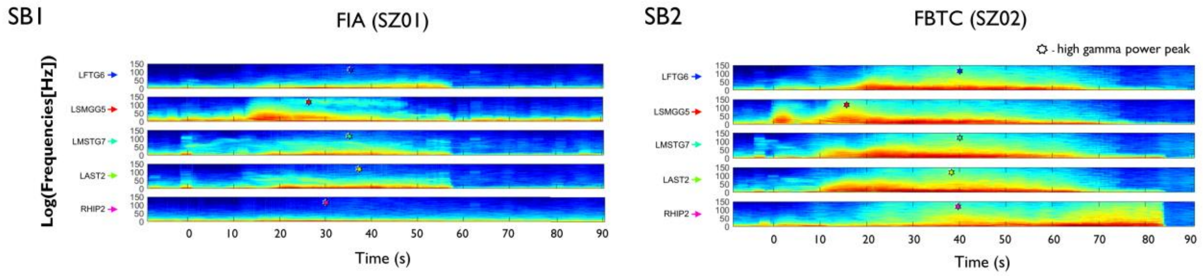


**Fig. S1**. Comparison of time-frequency representation for one FIA (SB1) and one FBTC (SB2) seizure displaying the (a)synchrony in high gamma power peaks, marked with colored dots on each representative channel.

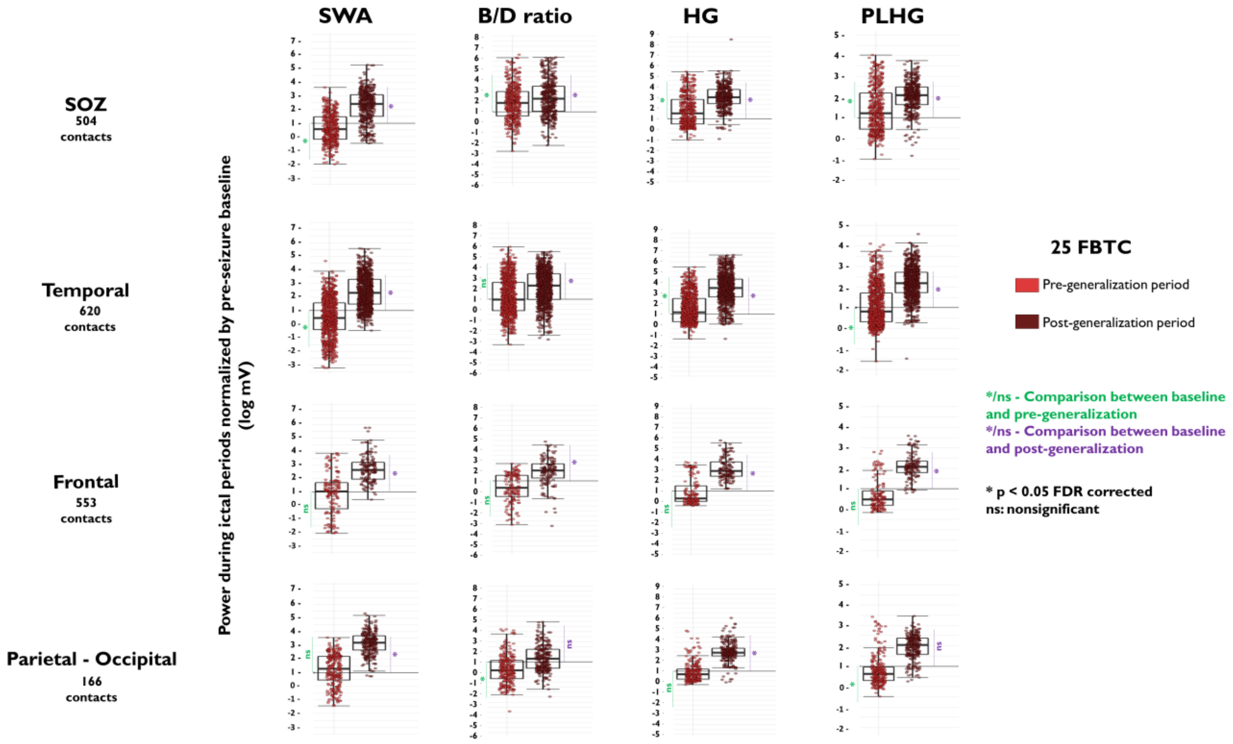


**Fig. S2**. Group results of the power spectra analysis for FBTC split by brain region (SOZ, temporal, frontal and parieto-occipital) and ictal period (pre- and post-generalization during FBTC) for SWA power, B/D, HG and PLHG. Each dot represents the normalized power value (normalized by baseline activity) for an electrode contact within each region. In this figure, pre-and post-generalization of FBTC are compared to pre-ictal baseline. Note that the values for baseline (pre-ictal) periods always equate to 1 since it is normalized by itself.

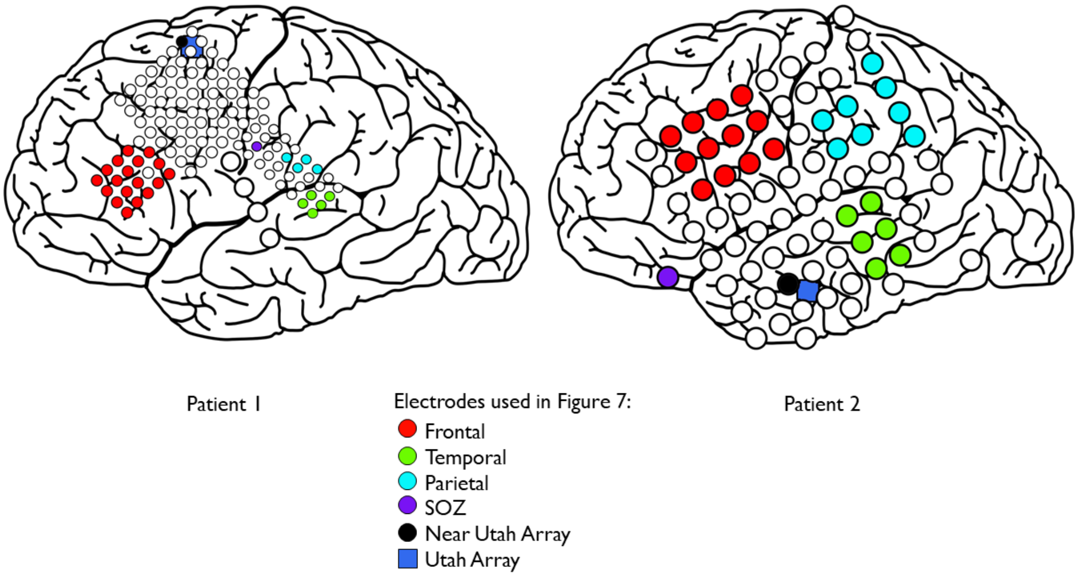


**Fig. S3**. Electrode maps for patients recorded with Utah arrays during FBTC.

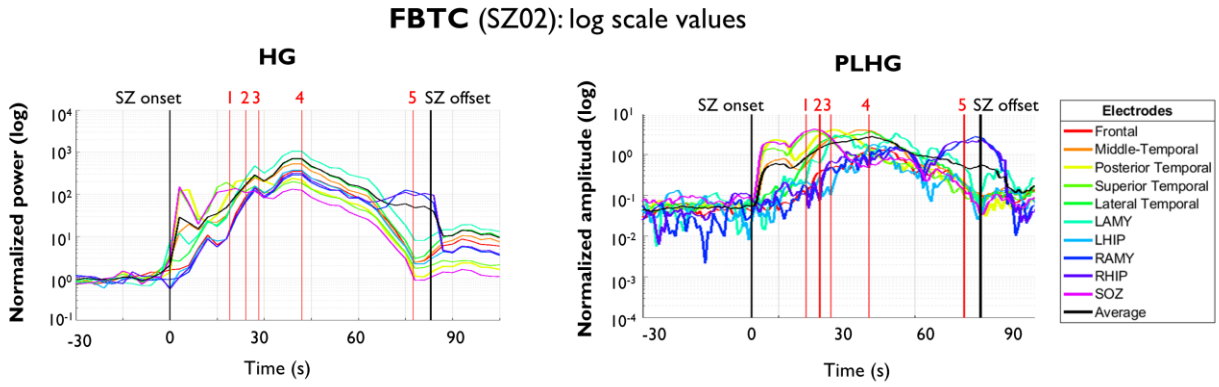


**Fig. S4**. Group results for High Gamma only in channels that did not pass the ER threshold during both FIA and FBTC, split by brain region (SOZ, temporal, frontal and parieto-occipital) and ictal period (first and second half of seizures). Each dot represents the normalized power value (normalized by baseline activity) for an electrode contact. FIA are displayed in blue and FBTC in red, with lighter colors indicating the first ictal period, and darker colors indicating the second ictal period. Black horizontal lines indicate values of pre-ictal baseline activity. These results suggest that HG increase in SOZ and temporal lobe but decreases in the rest of the brain during FIA. In contrast, HG diffusely increase starting at the onset of FBTC, and further build up as FBTC progress. HG was significantly higher during FBTC than during FIA for all brain areas and all ictal periods.
