## Supplementary material for "Distinct signatures of loss of consciousness during Focal Impaired Awareness versus Focal to Bilateral Tonic Clonic seizures": SupplMat_tables

### SUPPLEMENTARY MATERIALS

#### Tables

**Table S1a:** Descriptive statistics (mean ± SEM) of normalized SWA power values by brain region, seizure type and ictal period.

| | FIA | | FBTC | |
|---|---|---|---|---|
| | **First half** | **Second half** | **First half** | **Second half** |
| **SOZ** | 1.01 ± 0.03 | 1.67 ± 0.03 | 1.30 ± 0.05 | 2.77 ± 0.06 |
| **Temporal** | 1.04 ± 0.02 | 1.80 ± 0.02 | 1.44 ± 0.33 | 2.94 ± 0.04 |
| **Frontal** | 1.10 ± 0.04 | 1.61 ± 0.03 | 1.40 ± 0.05 | 2.92 ± 0.04 |
| **Parieto-Occipital** | 1.03 ± 0.03 | 1.50 ± 0.03 | 1.05 ± 0.06 | 2.63 ± 0.05 |

Note that the values for baseline (preictal) periods always equate to 1, since it is normalized by itself.

**Table S1b:** Descriptive statistics (mean ± SEM) of normalized BD power values by brain region, seizure type and ictal period.

| | FIA | | FBTC | |
|---|---|---|---|---|
| | **First half** | **Second half** | **First half** | **Second half** |
| **SOZ** | 1.46 ± 0.04 | 1.20 ± 0.04 | 2.37 ± 0.07 | 1.63 ± 0.08 |
| **Temporal** | 0.86 ± 0.03 | 0.70 ± 0.03 | 1.84 ± 0.05 | 1.55 ± 0.04 |
| **Frontal** | -0.47 ± 0.03 | -0.93 ± 0.03 | 1.39 ± 0.06 | 1.46 ± 0.06 |
| **Parieto-Occipital** | 0.25 ± 0.04 | -0.16 ± 0.05 | 1.96 ± 0.08 | 1.81 ± 0.07 |

Note that the values for baseline (preictal) periods always equate to 1, since it is normalized by itself.

**Table S1c:** Statistical comparison (*Z* values, (*p* values)) of normalized SWA power within seizure type, split by brain region and ictal period

| | FIA | | | FBTC | | |
|---|---|---|---|---|---|---|
| | **First half – Baseline** | **Second half – Baseline** | **Second half – First half** | **First half – Baseline** | **Second half – Baseline** | **Second half – First half** |

| | | | | | | |
|---|---|---|---|---|---|---|
| **SOZ** | 0.650 (1.00) | **23.003 (<0.001)** | **22.353 (< 0.001)** | **6.494 (< 0.001)** | **38.983 (<0.001)** | **32.489 (< 0.001)** |
| **Temporal** | 2.540 (0.68) | **41.529 (<0.001)** | **38.988 (< 0.001)** | **11.971 (< 0.001)** | **69.949 (<0.001)** | **57.978 (< 0.001)** |
| **Frontal** | 3.866 (0.02) | **24.054 (<0.001)** | **20.188 (< 0.001)** | **9.180 (< 0.001)** | **43.208 (<0.001)** | **34.028 (< 0.001)** |
| **Parieto-Occipital** | 0.936 (1.00) | **14.948 (<0.001)** | **14.012 (< 0.001)** | 1.236 (1.00) | **34.000 (<0.001)** | **32.765 (< 0.001)** |

Bold text indicates significant results (*p*<0.05). P values are corrected for multiple comparison using FDR correction.

**Table S1d:** Statistical comparison (*Z* values, (*p* values)) of normalized BD power within seizure type, split by brain region and ictal period.

| | **FIA** | | | **FBTC** | | |
|---|---|---|---|---|---|---|
| | **First half – Baseline** | **Second half – Baseline** | **Second half – First half** | **First half – Baseline** | **Second half – Baseline** | **Second half – First half** |
| **SOZ** | **11.309 (< 0.001)** | **4.837 (<0.001)** | **6.473 (< 0.001)** | **23.256 (< 0.001)** | **10.723 (< 0.001)** | **12.534 (< 0.001)** |
| **Temporal** | **-5.045 (< 0.001)** | **11.073 (<0.001)** | **6.028 (< 0.001)** | **23.116 (< 0.001)** | **15.934 (< 0.001)** | **6.520 (< 0.001)** |
| **Frontal** | **-41.510 (< 0.001)** | **54.391 (<0.001)** | **-12.882 (< 0.001)** | **6.256 (< 0.001)** | **7.435 (< 0.001)** | -1.179 (1.00) |
| **Parieto-Occipital** | **-16.107 (< 0.001)** | **30.819 (<0.001)** | **-8.823 (< 0.001)** | **12.615 (< 0.001)** | **11.119 (< 0.001)** | -1.496 (1.00) |

Bold text indicates significant results (*p*<0.05). P values are corrected for multiple comparison using FDR correction.

**Table S1e:** Statistical comparison (*Z* values, (*p* values)) of normalized SWA power by seizure type, brain region and ictal period.

| | | | FBTC | |
|---|---|---|---|---|
| | | | **First half** | **Second half** |
| **FIA** | **SOZ** | **First Half** | 1.126 (1.00) | 4.573 (<0.001) |
| | | **Second Half** | 6.591 (<0.001) | 3.144 (0.089) |
| | **Temporal** | **First Half** | 1.380 (1.00) | 7.146 (<0.001) |
| | | **Second Half** | 5.361 (<0.001) | 3.165 (0.085) |
| | **Frontal** | **First Half** | 1.449 (1.00) | 6.444 (<0.001) |
| | | **Second Half** | 4.121 (0.003) | 3.772 (0.012) |
| | **Parieto-Occipital** | **First Half** | 1.156 (1.00) | 7.029 (<0.001) |
| | | **Second Half** | 3.574 (0.023) | **4.611 (0.001)** |

Bold text indicates significant results (*p*<0.05). P values are corrected for multiple comparison using FDR correction.
Grey text indicates comparisons of less importance, which are also not shown in the figures.

**Table S1f:** Statistical comparison (*Z* values, (*p* values)) of normalized BD power by seizure type, brain region and ictal period.

| | | | FBTC | |
|---|---|---|---|---|
| | | | **First half** | **Second half** |
| **FIA** | **SOZ** | **First Half** | **6.723 (<0.001)** | 2.825 (0.203) |
| | | **Second Half** | 8.029 (<0.001) | 4.132 (0.002) |
| | **Temporal** | **First Half** | **7.195 (<0.001)** | 5.792 (<0.001) |
| | | **Second Half** | 8.013 (<0.001) | **6.610 (<0.001)** |
| | **Frontal** | **First Half** | **7.369 (<0.001)** | 7.726 (<0.001) |
| | | **Second Half** | 9.600 (<0.001) | **9.958 (<0.001)** |
| | **Parieto-Occipital** | **First Half** | **8.454 (<0.001)** | 7.968 (<0.001) |
| | | **Second Half** | 10.438 (<0.001) | **9.951 (<0.001)** |

Bold text indicates significant results (*p*<0.05). P values are corrected for multiple comparison using FDR correction.
Grey text indicates comparisons of less importance, which are also not shown in the figures.

**Table S2a:** Descriptive statistics (mean ± SEM) of normalized HG power values by brain region, seizure type and ictal period.

| | FIA | | FBTC | |
|---|---|---|---|---|
| | **First half** | **Second half** | **First half** | **Second half** |
| **SOZ** | 1.77 ± 0.04 | 1.85 ± 0.03 | 2.85 ± 0.06 | 3.75 ± 0.05 |
| **Temporal** | 1.79 ± 0.02 | 1.85 ± 0.02 | 3.31 ± 0.04 | 4.19 ± 0.04 |
| **Frontal** | 0.83 ± 0.02 | 1.04 ± 0.02 | 2.14 ± 0.05 | 3.19 ± 0.05 |
| **Parieto-Occipital** | 1.27 ± 0.04 | 1.39 ± 0.04 | 3.10 ± 0.06 | 4.06 ± 0.06 |

Note that the values for baseline (pre-ictal) periods always equate to 1, since it is normalized by itself.

**Table S2b:** Descriptive statistics (mean ± SEM) of normalized PLHG power values by brain region, seizure type and ictal period.

| | FIA | | FBTC | |
|---|---|---|---|---|
| | **First half** | **Second half** | **First half** | **Second half** |
| **SOZ** | 1.29± 0.02 | 1.35± 0.02 | 1.87 ± 0.03 | 2.24 ± 0.03 |
| **Temporal** | 1.17± 0.01 | 1.30 ± 0.01 | 1.89 ± 0.02 | 2.43 ± 0.02 |
| **Frontal** | 0.45 ± 0.01 | 0.60 ± 0.01 | 1.54 ± 0.03 | 2.01 ± 0.04 |
| **Parieto-Occipital** | 0.78 ± 0.02 | 0.86 ± 0.02 | 1.93± 0.03 | 2.52 ± 0.03 |

Note that the values for baseline (pre-ictal) periods always equate to 1, since it is normalized by itself.

**Table S2c:** Statistical comparison (*Z* values, (*p* values)) of normalized HG power within seizure type, split by brain region and ictal period.

| | FIA | | | FBTC | | |
|---|---|---|---|---|---|---|
| | **First half – Baseline** | **Second half – Baseline** | **Second half – First half** | **First half – Baseline** | **Second half – Baseline** | **Second half – First half** |

| | | | | | | |
|---|---|---|---|---|---|---|
| **SOZ** | **24.842 (< 0.001)** | **27.103 (< 0.001)** | 2.261 (0.843) | **38.370 (< 0.001)** | **56.885 (< 0.001)** | **18.515 (< 0.001)** |
| **Temporal** | **38.375 (< 0.001)** | **41.697 (< 0.001)** | 3.322 (0.097) | **78.263 (< 0.001)** | **107.766 (< 0.001)** | **29.503 (< 0.001)** |
| **Frontal** | **-6.061 (< 0.001)** | 1.705 (0.98) | **7.766 (< 0.001)** | **24.271 (< 0.001)** | **46.311 (< 0.001)** | **22.040 (< 0.001)** |
| **Parieto-Occipital** | **-7.858 (< 0.001)** | **10.948 (< 0.001)** | 3.090 (0.191) | **41.226 (< 0.001)** | **60.004 (< 0.001)** | **18.777 (< 0.001)** |

Bold text indicates significant results ($p$<0.05). P values are corrected for multiple comparison using FDR correction.

**Table S2d:** Statistical comparison ($Z$ values, ($p$ values)) of normalized PLHG power within seizure type, split by brain region and ictal period.

| | **FIA** | | | **FBTC** | | |
|---|---|---|---|---|---|---|
| | **First half – Baseline** | **Second half – Baseline** | **Second half – First half** | **First half – Baseline** | **Second half – Baseline** | **Second half – First half** |
| **SOZ** | **14.648 (< 0.001)** | **17.680 (< 0.001)** | 3.032 (0.195) | **25.429** (< **0.001**) | **36.243 (< 0.001)** | **10.814 (< 0.001)** |
| **Temporal** | **13.508 (< 0.001)** | **23.560 (< 0.001)** | **10.051 (< 0.001)** | **45.519 (< 0.001)** | **73.392 (< 0.001)** | **27.872 (< 0.001)** |
| **Frontal** | **-31.140 (< 0.001)** | **-23.132 (< 0.001)** | **8.008 (< 0.001)** | **14.774 (< 0.001)** | **27.337 (< 0.001)** | **12.564 (< 0.001)** |
| **Parieto-Occipital** | **-9.491 (< 0.001)** | **-5.793 (< 0.001)** | 3.698 (0.024) | **26.783 (< 0.001)** | **43.681 (< 0.001)** | **16.898 (< 0.001)** |

Bold text indicates significant results ($p$<0.05). P values are corrected for multiple comparison using FDR correction.

**Table S2e:** Statistical comparison (*Z* values, (*p* values)) of normalized HG power by seizure type, brain region and ictal period.

| | | | FBTC | |
|---|---|---|---|---|
| | | | **First half** | **Second half** |
| **FIA** | **SOZ** | **First Half** | **5.389 (<0.001)** | 9.904 (<0.001) |
| | | **Second Half** | 5.031 (<0.001) | **9.546 (<0.001)** |
| | **Temporal** | **First Half** | **6.481 (<0.001)** | 10.941 (<0.001) |
| | | **Second Half** | 6.133 (<0.001) | **10.592 (<0.001)** |
| | **Frontal** | **First Half** | **6.623 (<0.001)** | 11.872 (<0.001) |
| | | **Second Half** | 5.568 (<0.001) | **8.831 (<0.001)** |
| | **Parieto-Occipital** | **First Half** | **7.162 (<0.001)** | 11.976 (<0.001) |
| | | **Second Half** | 6.614 (<0.001) | **11.429 (<0.001)** |

Bold text indicates significant results ($p$<0.05). P values are corrected for multiple comparison using FDR correction.
Grey text indicates comparisons of less importance, which are also not shown in the figures.

**Table S2f:** Statistical comparison (*Z* values, (*p* values)) of normalized PLHG power by seizure type, brain region and ictal period.

| | | | FBTC | |
|---|---|---|---|---|
| | | | **First half** | **Second half** |
| **FIA** | **SOZ** | **First Half** | **8.267 (<0.001)** | 17.661 (<0.001) |
| | | **Second Half** | 7.409 (<0.001) | **12.658 (<0.001)** |
| | **Temporal** | **First Half** | **10.136 (<0.001)** | 18.241 (<0.001) |
| | | **Second Half** | 8.172 (<0.001) | **16.277 (<0.001)** |
| | **Frontal** | **First Half** | **12.738 (<0.001)** | 19.209 (<0.001) |
| | | **Second Half** | 10.814 (<0.001) | **17.285 (<0.001)** |
| | **Parieto-Occipital** | **First Half** | **13.465 (<0.001)** | 21.705 (<0.001) |
| | | **Second Half** | 12.302 (<0.001) | **20.542 (<0.001)** |

Bold text indicates significant results ($p$<0.05). P values are corrected for multiple comparison using FDR correction.
Grey text indicates comparisons of less importance, which are also not shown in the figures.

**Table S3: (**A)synchrony in the recruitment of channels measured with Epileptogenicity Index (E)I), delta power and the amplitude of the slow waves for CPS and GTC seizures.

| Parameter vs all channels | FIA | | FBTC | | Statistics (p-value, Cohen's d) |
|---|---|---|---|---|---|
| ER passed channels | 45,6% | | 69.1% | | p < 0.001; d = 0.49 |
| ER passed channels – first vs second half of the seizure | 36.4% (79.9%) | 9.2% (20.1%) | 48.7% (70.5%) | 20.4% (28.8%) | Between seizure types: p<0.001 d =0.25(first halves), d = 0.32 (second halves)<br>Within: p<0.001; d = 0.70 (CPS), d = 0.34 (GTC) |
| (a)synchrony of ER passed | 10% | | 23% | | p < 0.001; d = 0.84 |
| (a)synchrony of ER passed vs channels that passed ER | 29% | | 35% | | p = 0.044; d = 0.20 |
| (a)synchrony of delta power (10 sec bins, 1 sec apart) | 28% | | 34% | | p = 0.025; d = 0.33 |
| (a)synchrony of SWs maximal negative peak (WindowSize=10, Step=1 s) | 35% | | 42% | | p = 0.012; d = 0.67 |

Bold text indicates significant results (*p*<0.001). P values are corrected for multiple comparison using FDR correction.

**Table S4:** Differences in the recruitment of individual channels measured with EI in various brain areas between the first and the second half of the seizure for FIA and FBTC and between pre-generalized and post-generalized period for FBTC seizures.

| | FIA | | | | FBTC | | | | | | |
|---|---|---|---|---|---|---|---|---|---|---|---|
| **Channels ratios/ brain regions** | Passed ER vs all | First Half vs all passed | Second Half vs all passed | First vs Second half (p values) | Passed ER vs all | First Half vs all passed | Second Half vs all passed | First vs Second half (p values) | Pre-gen vs all passed | Post-gen Half vs all passed | Pre-gen vs Post-gen (p values) |
| **SOZ** | 60.2% | 54.9% (91.3%) | 5.7% (9.7%) | p < 0.001 | 69.1% | 61.8% (89.4%) | 7.4% (10.6%) | p < 0.001 | 81.2% | 18.8% | p < 0.001 |
| **Temporal** | 47.2% | 37.7% (79.8%) | 9.5% (20.2%) | p < 0.001 | 69.3% | 51.9% (74.2%) | 18.0% (25.8%) | p < 0.001 | 59.9% | 40.1% | p < 0.001 |

| | | | | | | | | | | | |
|---|---|---|---|---|---|---|---|---|---|---|---|
| **Temporal posterior lateral** | 35.2% | 26.5% (75.2%) | 10.5% (24.8%) | p < 0.001 | 78.8% | 53.9% (68.%) | 23.7% (31.3%) | p < 0.001 | 35.0% (59.6%) | 23.7% (40.4%) | p < 0.001 |
| **Frontal** | 48.6% | 38.1% (78.5%) | 10.5% (21%) | p < 0.001 | 62.7% | 36.8% (58.7%) | 25.9% (41.3%) | p = 0.019 | 41.0% | 59.0% | p = 0.010 |
| **Prefrontal** | 49.8% | 38.7% (77,8%) | 11.1% (22,2%) | p < 0.001 | 62.2% | 37.6% (60.4%) | 24.6% (39.6%) | p = 0.023 | 41.4% | 58.6% | p = 0.017 |
| **Motor** | 43.7% | 35.9% (82.2%) | 7.8% (17.8%) | p < 0.001 | 64.3% | 34.3% (53.3%) | 30.1% (46.7%) | p = 0.260 | 39.8% | 60.2% | p < 0.001 |
| **Parietal** | 29.3% | 23.7% (80.9%) | 5.6% (19.1%) | p < 0.001 | 73.6% | 53.1% (72.2%) | 20.5% (27.8%) | p < 0.001 | 64.9% | 35.7% | p < 0.001 |
| **Occipital** | 39.4% | 34.5% (87.4%) | 5.0% (12.6%) | p < 0.001 | 78.3% | 55.5% (70.9%) | 22.8% (29.1%) | p < 0.001 | 61.5% | 38.5% | p < 0.001 |
| **Limbic** | 51.1% | 42.0% (82.4%) | 9.0% (17.6%) | p < 0.001 | 58.8% | 45.4% (75.5%) | 14.4% (24.5%) | p < 0.001 | 64.3% | 35.7% | p < 0.001 |

Bold text indicates significant results (*p*<0.001). P values are corrected for multiple comparison using FDR correction.

**Table S5:** Comparison between the recruitment of channels measured with EI between first and the second half in FIA and FBTC, and pre-generalization and post-generalization periods for FBTC (*p* values).

| | | **FBTC** | | | |
|---|---|---|---|---|---|
| | | **Pre-generalization** | **Post-generalization** | **First Half** | **Second Half** |
| **SOZ** | **First Half** | p = 0.067 | p < 0.001 | p = 0.010 | p < 0.001 |
| | **Second Half** | p < 0.001 | p < 0.001 | p < 0.001 | p = 0.028 |
| **Temporal** | **First Half** | p = 0.023 | p = 0.010 | p < 0.001 | p < 0.001 |
| | **Second Half** | p < 0.001 | p < 0.001 | p < 0.001 | p < 0.001 |
| **Temporal posterior lateral** | **First Half** | | | | |
| | **Second Half** | | | | |
| **Frontal** | **First Half** | p < 0.001 | p = 0.781 | p = 0.340 | p = 0.012 |
| | **Second Half** | p < 0.001 | p < 0.001 | p < 0.001 | p < 0.001 |
| | **First Half** | p = 0.423 | p = 0.012 | p < 0.001 | p = 0.212 |

| | | | | | | |
|---|---|---|---|---|---|---|
| **FIA** | **Frontal - Prefrontal** | **Second Half** | p < 0.001 | p < 0.001 | p < 0.001 | p < 0.001 |
| | **Frontal - Motor** | **First Half** | p < 0.001 | p = 0.198 | p = 0.352 | p = 0.109 |
| | | **Second Half** | p < 0.001 | p < 0.001 | p < 0.001 | p < 0.001 |
| | **Parietal** | **First Half** | p < 0.001 | p = 0.021 | p < 0.001 | p = 0.064 |
| | | **Second Half** | p < 0.001 | p < 0.001 | p < 0.001 | p < 0.001 |
| | **Occipital** | **First Half** | p < 0.001 | p = 0.050 | p < 0.001 | p = 0.009 |
| | | **Second Half** | p < 0.001 | p < 0.001 | p < 0.001 | p < 0.001 |
| | **Limbic** | **First Half** | p = 0.192 | p < 0.001 | p = 0.213 | p < 0.001 |
| | | **Second Half** | p < 0.001 | p < 0.001 | p < 0.001 | p = 0.021 |

Bold text indicates significant results (*p*<0.001). P values are corrected for multiple comparison using FDR correction. Grey text indicates comparisons of less importance, which are also not shown in the figures.

**Table S6a**: Descriptive statistics (mean ± SEM) of the percentage of explained variance in HG for EMG, iEEG, depth iEEG and superficial iEEG electrodes.

| | **Baseline** | **Pre-generalization** | **Post-generalization** |
|---|---|---|---|
| **EMG** | 0.253 ± 0.098 | 2.311 ± 0.876 | 1.364 ± 0.427 |
| **iEEG** | 13.013 ± 3.455 | 54.884 ± 7.222 | 61.264 ± 5.528 |
| **Deep (iEEG)** | 25.868 ± 7.531 | 64.702 ± 8.039 | 77.928 ± 4.782 |
| **Superficial (iEEG)** | 16.196 ± 5.998 | 48.982 ± 7.604 | 61.120 ± 5.315 |

Note: In seizures for which the point of behavioral generalization was not available from behavioral data, the point of generalization was estimated based on the average delay from the maximum slope increase in PLHG power (cf. Fig. 4D and Table S5).

**Table S6b**: Comparisons between EMG and iEEG values, expressed as t values (p values).

| | | **iEEG** | | |
|---|---|---|---|---|
| | | **Baseline** | **Pre-generalization** | **Post-generalization** |
| **EMG** | **Baseline** | **3.69 (0.002)** | **7.563 (<0.001)** | **11.034 (<0.001)** |
| | **Pre-generalization** | **3.002 (0.007)** | **9.4 (< 0.001)** | **10.532 (<0.001)** |
| | **Post-generalization** | **3.345 (0.003)** | **7.397 (<0.001)** | **10.71 (< 0.001)** |

Bold text indicates significant results (*p*<0.05). P values are FDR corrected for multiple comparison.

**Table S6c**: Comparisons between deep iEEG and superficial iEEG values, expressed as t values (p values).

| | | **Superficial (iEEG)** | | |
|---|---|---|---|---|
| | | **Baseline** | **Pre-generalization** | **Post-generalization** |
| | **Baseline** | - 1.004 (0.323) | 1.914 (0.064) | **3.627 (< 0.001)** |
| **Deep** | **Pre-generalization** | **- 4.517 (<0.001)** | - 1.364 (0.182) | - 0.309 (0.759) |
| **(iEEG)** | **Post-generalization** | **-8.043 (<0.001)** | **- 3.321 (0.002)** | **- 2.513 (0.017)** |

Bold text indicates significant results (*p*<0.05). P values are FDR corrected for multiple comparison. Wherever it was available, depth electrodes implanted into *Amygdala* were used.

**Table S6d**: Comparisons between deep iEEG and superficial iEEG values, expressed as t values (p values).

| | | **Superficial (iEEG)** | | |
|---|---|---|---|---|
| | | **Baseline** | **Pre-generalization** | **Post-generalization** |
| | **Baseline** | -1.376 (0.741) | 2.409 (0.165) | **3.532 (0.008)** |
| **Deep** | **Pre-generalization** | **-4.882 (<0.001)** | -1.096 (0.881) | 0.027 (1.00) |
| **(iEEG)** | **Post-generalization** | **-5.312 (<0.001)** | 1.526 (0.648) | -0.403 (0.998) |

Bold text indicates significant results (*p*<0.05). P values are FDR corrected for multiple comparison. Wherever it was available, depth electrodes implanted into *Hippocampus* were used.

**Table S7a:** Statistical comparison (*Z* values, (*p* values)) of normalized SWA power for FBTC, contrasting brain regions and ictal periods (25 FBTC).

| | FBTC | | |
|---|---|---|---|
| | **Pre-generalization – Baseline** | **Post-generalization – Baseline** | **Post-generalization – Pre-generalization** |
| **SOZ** | **6.594 (<0.001)** | **-23.355 (<0.001)** | **-29.859 (<0.001)** |
| **Temporal** | **13.806 (<0.001)** | **-36.448 (<0.001)** | **-50.190 (<0.001)** |
| **Frontal** | 1.623 (1.00) | **-18.138 (<0.001)** | **-19.752 (<0.001)** |
| **Parieto-Occipital** | -2.768 (0.230) | **-30.008 (<0.001)** | **-27.241 (<0.001)** |

Bold text indicates significant results ($p<0.05$). P values are corrected for multiple comparison using FDR correction.

**Table S7b:** Statistical comparison (*Z* values, (*p* values)) of normalized B/D ratio power for FBTC, contrasting brain regions and ictal periods (25 FBTC).

| | FBTC | | |
|---|---|---|---|
| | **Pre-generalization – Baseline** | **Post-generalization – Baseline** | **Post-generalization – Pre-generalization** |
| **SOZ** | **-12.017 (<0.001)** | **-17.967 (<0.001)** | **-6.114 (<0.001)** |
| **Temporal** | **-4.082 (0.003)** | **-21.954 (<0.001)** | **-17.891 (<0.001)** |
| **Frontal** | **5.372 (<0.001)** | **-6.486 (<0.001)** | **-11.829 (<0.001)** |
| **Parieto-Occipital** | **7.685 (<0.001)** | **-3.440 (0.033)** | **-11.125 (<0.001)** |

Bold text indicates significant results ($p<0.05$). P values are corrected for multiple comparison using FDR correction.

**S7c:** Average temporal distance (mean ± SEM, in sec) from the peak in power of Delta and B/D ratio

| | **Δ time from Seizure Start** | **Δ time from Behavioral Generalization** | **Δ time from Seizure End** |
|---|---|---|---|
| **Delta** | -100.96 ± 16.41 | 37 ± 8.15 | 59.16 ± 11.01 |
| **B/D ratio** | -70.36 ± 14.66 | 22.92 ± 5.05 | 88.12 ± 12.17 |

**Table S8a:** Statistical comparison (*Z* values, (*p* values)) of normalized HG power for FBTC, contrasting brain regions and ictal periods (25 FBTC).

| | | | **FBTC** | |
|---|---|---|---|---|
| | | | **Pre-generalization** | **Post-generalization** |
| **FBTC** | **SOZ** | **Baseline** | **13.468 (<0.001)** | **37.331 (<0.001)** |
| | **Temporal** | **Baseline** | **12.852 (<0.001)** | **63.476 (<0.001)** |
| | **Frontal** | **Baseline** | -0.968 (1.00) | **21.685 (<0.001)** |
| | **Parieto-Occipital** | **Baseline** | -1.100 (1.00) | **24.475 (<0.001)** |

Bold text indicates significant results (*p*<0.05). P values are corrected for multiple comparison using FDR correction.

**Table S8b:** Statistical comparison (*Z* values, (*p* values)) of normalized PLHG power for FBTC, contrasting brain regions and ictal periods (25 FBTC).

| | | | **FBTC** | |
|---|---|---|---|---|
| | | | **Pre-generalization** | **Post-generalization** |
| **FBTC** | **SOZ** | **Baseline** | **9.406 (<0.001)** | **26.542 (<0.001)** |
| | **Temporal** | **Baseline** | -2.546 (0.749) | **43.224 (<0.001)** |
| | **Frontal** | **Baseline** | **- 4.125 (0.007)** | **15.883 (<0.001)** |
| | **Parieto-Occipital** | **Baseline** | **-3.949 (0.016)** | **18.228 (<0.001)** |

Bold text indicates significant results (*p*<0.05). P values are corrected for multiple comparison using FDR correction.

**Table S8c:** Average temporal distance (mean ± SEM, in sec) from the peak in power of HG and PLHG.

| | **Δ time from Seizure Start** | **Δ time from Behavioral Generalization** | **Δ time from Seizure End** |
|---|---|---|---|
| **HG** | -72 ± 13.2 | 21 ± 3.7 | 87.3 ± 12.2 |
| **PLHG** | -71 ± 12.86 | 85.6 ± 11.9 | 88.12 |

**Table S9a:** Statistical comparison (*Z* values, (*p* values)) of normalized SWA power between seizure type, brain region and ictal period (25 FBTC; 57 FIA).

| | | | **FBTC** | |
|---|---|---|---|---|
| | | | **Pre-generalization** | **Post-generalization** |
| | **SOZ** | **First Half** | -1.312 (1.00) | 6.063 (0.01) |
| | | **Second Half** | - 4.663(0.004) | 2.717 (0.278) |
| | **Temporal** | **First Half** | - 3.134 (0.094) | 5.685 (<0.001) |
| **FIA** | | **Second Half** | - 6.311 (<0.001) | 2.508 (0.421) |
| | **Frontal** | **First Half** | - 3.134 (0.094) | 5.883 (<0.001) |
| | | **Second Half** | - 4.753 (<0.001) | 0.462 (1.00) |
| | **Parieto-Occipital** | **First Half** | - 1.284 (1.00) | 7.173(<0.001) |
| | | **Second Half** | - 3.497(0.037) | **4.960 (<0.001)** |

Bold text indicates significant results (*p*<0.05). P values are corrected for multiple comparison using FDR correction.
Grey text indicates comparisons of less importance, which are also not shown in the figures.

**Table S9b:** Statistical comparison (*Z* values, (*p* values)) of normalized B/D ratio power between seizure type, brain region and ictal period (25 FBTC; 57 FIA).

| | | | **FBTC** | |
|---|---|---|---|---|
| | | | **Pre-generalization** | **Post-generalization** |
| | **SOZ** | **First Half** | **3.919 (0.006)** | 6.728 (<0.001) |
| | | **Second Half** | 3.979 (0.005) | **6.788 (<0.001)** |
| | **Temporal** | **First Half** | 3.253 (0.061) | 8.483 (<0.001) |
| **FIA** | | **Second Half** | 4.431(<0.001) | **9.660 (<0.001)** |

| | | | |
|---|---|---|---|
| **Frontal** | **First Half** | **4.048 (0.003)** | 9.652 (<0.001) |
| | **Second Half** | 6.454 (<0.001) | **12.089 (<0.001)** |
| **Parieto-Occipital** | **First Half** | **4.108 (0.003)** | 8.703(<0.001) |
| | **Second Half** | 7.019 (<0.001) | **11.614 (<0.001)** |

Bold text indicates significant results (*p*<0.05). P values are corrected for multiple comparison using FDR correction.
Grey text indicates comparisons of less importance, which are also not shown in the figures.

**Table S10a:** Statistical comparison (*Z* values, (*p* values)) of normalized HG power between seizure type, brain region and ictal period (25 FBTC; 57 FIA).

| | | | **FBTC** | |
|---|---|---|---|---|
| | | | **Pre-generalization** | **Post-generalization** |
| | **SOZ** | **First Half** | 3.176 (0.096) | 9.840 (<0.001) |
| | | **Second Half** | 1.167 (1.00) | **7.834 (<0.001)** |
| | **Temporal** | **First Half** | 0.519 (1.00) | 10.618 (<0.001) |
| **FIA** | | **Second Half** | 0.667(1.00) | **9.432 (<0.001)** |
| | **Frontal** | **First Half** | 1.425 (1.00) | 9.541 (<0.001) |
| | | **Second Half** | 0.547 (1.00) | **8.663 (<0.001)** |
| | **Parieto-Occipital** | **First Half** | 2.361 (0.624) | 10.697 (<0.001) |
| | | **Second Half** | 0.898(1.00) | **9.234 (<0.001)** |

Bold text indicates significant results (*p*<0.05). P values are corrected for multiple comparison using FDR correction.
Grey text indicates comparisons of less importance, which are also not shown in the figures.

**Table S10b:** Statistical comparison (*Z* values, (*p* values)) of normalized PLHG power between seizure type, brain region and ictal period (25 FBTC; 57 FIA).

| | | **FBTC** | |
|---|---|---|---|
| | | **Pre-generalization** | **Post-generalization** |
| **SOZ** | **First Half** | **5.187 (<0.001)** | 10.702 (<0.001) |
| | **Second Half** | 3.120 (0.100) | **8.680 (<0.001)** |
| **Temporal** | **First Half** | 1.346 (1.00) | 12.318 (<0.001) |

| | | | | |
|---|---|---|---|---|
| **FIA** | | **Second Half** | 0.266 (1.00) | **10.707 (<0.001)** |
| | **Frontal** | **First Half** | 2.009 (0.868) | 12.087 (<0.001) |
| | | **Second Half** | 0.811 (1.00) | **10.889 (<0.001)** |
| | **Parieto-Occipital** | **First Half** | 3.899 (0.007) | 13.343 (<0.001) |
| | | **Second Half** | 2.794 (0.237) | **12.239 (<0.001)** |

Bold text indicates significant results (*p*<0.001). P values are corrected for multiple comparison using FDR correction.
Grey text indicates comparisons of less importance, which are also not shown in the figures.

**Table S11a:** Descriptive statistics (mean ± SEM) of normalized HG power by brain region, seizure type and ictal period, only for channels that did not pass the ER threshold.

| | **FIA** | | **FBTC** | |
|---|---|---|---|---|
| | **First half** | **Second half** | **First half** | **Second half** |
| **SOZ** | 1.47 ± 0.05 | 1.62 ± 0.04 | 3.37 ± 0.11 | 4.19± 0.08 |
| **Temporal** | 1.55 ± 0.03 | 1.71± 0.03 | 3.67± 0.07 | 4.35 ± 0.07 |
| **Frontal** | 0.72 ± 0.04 | 0.99 ± 0.03 | 2.54 ± 0.17 | 3.21 ± 0.17 |
| **Parieto-Occipital** | 1.05 ± 0.04 | 1.25 ± 0.04 | 4.11 ± 0.14 | 5.06 ± 0.14 |

Note that the values for baseline (pre-ictal) periods always equate to 1, since it is normalized by itself.

**Table S11b:** Statistical comparison (*Z* values, (*p* values)) of normalized HG power within seizure type, split by brain region and ictal period, only for channels that did not pass the ER threshold.

| | **FIA** | | | **FBTC** | | |
|---|---|---|---|---|---|---|
| | **First half – Baseline** | **Second half – Baseline** | **Second half – First half** | **First half – Baseline** | **Second half – Baseline** | **Second half – First half** |
| **SOZ** | **11.572 (< 0.001)** | **15.038 (< 0.001)** | 3.466 (0.067) | **24.660 (< 0.001)** | **34.196 (< 0.001)** | **9.073 (< 0.001)** |
| **Temporal** | **19.737 (< 0.001)** | **25.301 (< 0.001)** | **5.564 (<0.001)** | **47.953 (< 0.001)** | **60.802 (< 0.001)** | **12.565 (< 0.001)** |

| | | | | | | |
|---|---|---|---|---|---|---|
| **Frontal** | **-6.854 (< 0.001)** | 0.245 (1.00) | **6.609 (< 0.001)** | **13.607 (< 0.001)** | **20.105 (< 0.001)** | **22.040 (< 0.001)** |
| **Parieto-Occipital** | -1.338 (1.00) | **-6.597 (< 0.001)** | **-5.259 (<0.001)** | **26.365 (< 0.001)** | **34.500 (< 0.001)** | **6.152 (< 0.001)** |

Bold text indicates significant results ($p$<0.05). P values are corrected for multiple comparison using FDR correction.
Grey text indicates comparisons of less importance, which are also not shown in the figures.

**Table S11c:** Statistical comparison ($Z$ values, ($p$ values)) of normalized HG power by seizure type, brain region and ictal period, only for channels that did not pass the ER threshold.

| | | | **FBTC** | |
|---|---|---|---|---|
| | | | **First half** | **Second half** |
| | **SOZ** | **First Half** | **6.195 (<0.001)** | 10.220 (<0.001) |
| | | **Second Half** | 5.554 (<0.001) | **9.579 (<0.001)** |
| | **Temporal** | **First Half** | **7.485 (<0.001)** | 10.789 (<0.001) |
| **FIA** | | **Second Half** | 6.754 (<0.001) | **10.057 (<0.001)** |
| | **Frontal** | **First Half** | **7.727 (<0.001)** | 10.926 (<0.001) |
| | | **Second Half** | 6.580 (<0.001) | **9.774 (<0.001)** |
| | **Parieto-Occipital** | **First Half** | **8.041 (<0.001)** | 12.208 (<0.001) |
| | | **Second Half** | 7.026 (<0.001) | **11.192 (<0.001)** |

Bold text indicates significant results ($p$<0.05). P values are corrected for multiple comparison using FDR correction.
Grey text indicates comparisons of less importance, which are also not shown in the figures.
